# Maternal Vitamin C Deficiency and Genetic Risk Factors Contribute to Congenital Defects through Dysregulation of DNA Methylation

**DOI:** 10.1101/2025.05.27.656260

**Authors:** Bernard K. van der Veer, Colin Custers, Wannes Brangers, Riet Cornelis, Spyridon Champeris Tsaniras, Kobe De Ridder, Bernard Thienpont, Huiyong Cheng, Qiuying Chen, Daniel Kraushaar, Richard H. Finnell, Steven S. Gross, Kian Peng Koh

**Affiliations:** Laboratory for Stem Cell and Developmental Epigenetics, Department of Development and Regeneration, KU Leuven, Leuven 3000, Belgium; Laboratory for Functional Epigenetics, Department of Human Genetics, KU Leuven, Leuven 3000, Belgium; Department of Pharmacology, Weill Cornell Medical College, New York, NY 10065, USA; Department of Molecular and Cellular Biology, Genomic and RNA Profiling Core, Baylor College of Medicine, Houston, Texas, USA; Department of Molecular and Cellular Biology, Center for Precision Environmental Health, Baylor College of Medicine, Houston, Texas, USA; Department of Molecular and Human Genetics, Department of Medicine, Baylor College of Medicine, Houston, Texas, USA

## Abstract

Maternal dietary insufficiencies can reshape the fetal epigenome during gestation, contributing to birth defects and developmental disorders. Vitamin C (VitC) is a critical co-factor for Ten-Eleven-Translocation (TET) DNA demethylases, but the impact of its deficiency on embryonic development has gone largely unappreciated. Here, we show that maternal VitC deficiency in L-gulonolactone oxidase (*Gulo*)-deficient mice, which like humans are unable to synthesize VitC, can cause highly penetrant developmental delays and malformations in non-inbred embryos during the vulnerable period of gastrulation. DNA hypermethylation in *Gulo*^-/-^ embryonic neural tissues of susceptible strains increases with VitC dose reduction and with the severity of embryonic pathologies, coinciding with hallmarks of TET1 dysfunction. A moderate reduction in VitC status is sufficient to induce DNA hypermethylation and cause neural tube defects. Our results suggest that promoting timely VitC supplementation by at-risk pregnant mothers may prevent a range of birth defects and enhance health outcomes of future generations.

## Introduction

Pre-natal exposures to adverse environmental insults can contribute to a broad spectrum of fetal pathologies including birth defects and developmental disorders (*1*). The concept of a developmental origin of health and disease, first proposed by the British epidemiologist David Barker in 1990, now encompasses the definition of “a sensitive window of vulnerability” in early life, when environmental exposures can induce long-lasting disruption of developmental programming to perturb cellular functions later in life (*2*). One major source of environmental influence *in utero* is maternal nutritional status, the major contributor to birth defects in approximately 1 in 33 infants born in the United States and the leading cause of death for infants during the first year of life (*3*). Uncovering environmental vulnerabilities in early life creates opportunities for interventions. For example, food fortification programs to increase periconceptional intake of folic acid (FA, or vitamin B9) have reduced the prevalence of neural tube defects (NTDs), the second most common structural congenital disorder, by 30-50% (*4*). However, very few similar opportunities to reduce fetal pathologies have been identified, due in large part to significant knowledge gaps of the underlying mechanisms governing how early life environmental exposures interact with genetic risk factors to cause adverse developmental outcomes.

The epigenome consists of a layer of biochemical modifications of chromatin that regulates gene expression, thereby providing a molecular interface mediating environmental (including nutritional cofactors) interactions with the genome. In early mammalian development, the epigenome undergoes a highly dynamic remodeling that involves spatial and temporal coordination of changes in DNA methylation and histone post-translational modifications (PTM). Although erasure of PTMs can occur passively by evicting marked histones during DNA replication, more recently discovered histone and DNA demethylases are now known to participate actively in epigenomic reprogramming during early development. Of particular interest, the Jumonji domain C (JMJC) family of Fe^2+^ and α-ketoglutarate (αKG)-dependent dioxygenases include histone H3 lysine-specific demethylases (KDMs) and the Ten-Eleven-Translocation (TET) DNA demethylases (TET1, TET2 and TET3) (*5*). Histone lysine-specific demethylation occurs by oxidation of methyl groups and release of formaldehyde, whereas DNA demethylation involves iterative oxidation of 5-methylcytosine (5mC) into 5-hydroxymethylcytosine (5hmC) and further oxidation products (*6–8*). TET-mediated DNA demethylation at enhancers and promoters during peri-implantation regulates lineage fate decisions and primes the epigenome for organogenesis (*9–11*), while disruption of histone methylation reactions has been shown to adversely affect gastrulation (*12*). The dependency of TET and JmjC domain family dioxygenases on cellular redox (Fe^2+^) and metabolic substrates (αKG) also positions them as environmental sensors of nutritional inputs, such that their peak activities during early life epigenome remodeling may underlie the windows of the developmental period that is highly vulnerable to environmental exposures.

L-ascorbic acid, commonly known as vitamin C (VitC), is an essential nutrient that can have a profound impact on the epigenome by enhancing the demethylating activities of DNA and histone dioxygenases (*13*). Ascorbate, the dominant form of VitC under physiological pH conditions, maintains these dioxygenases in their fully active form by reducing catalytically inhibitory oxidized iron species (mostly Fe^3+^) and regenerating Fe^2+^ at their enzymatically active site(*13–15*). In cell-based and *in vitro* assays, ascorbate enhances TET catalytic activity directly to generate 5hmC, but treatment with other reducers such as glutathione does not, suggesting that VitC is not functioning solely as a general reducer but as a specific regulator of dioxygenases (*13*). VitC treatment of mouse embryonic stem cell (ESC) cultures results in rapid and global TET-dependent increase in 5hmC, followed by DNA demethylation at gene promoters (*16*). VitC also enhances the generation of induced pluripotent stem cells from terminally differentiated cells during transcription factor-driven reprogramming, by stimulating dioxygenases to remove epigenetic barriers imposed by histone and DNA methylation (*17, 18*). Nonetheless, evidence for the role of VitC in regulating dioxygenase functions during peri-gastrulation development *in vivo* is still lacking.

Most vertebrates and mammals, such as rodents, synthesize VitC *de novo* in the liver primarily from glucose, through a pathway whose final step is catalyzed by the enzyme L-gulonolactone oxidase (GULO) (*13, 19*). In contrast, humans and higher primates lack a functional form of GULO and rely on a supply of ascorbate entirely through the diet(*20*). A common model for VitC deficiency is the *Gulo*^tm1Mae^ mouse, which carries a targeted knockout (KO) mutation of the *Gulo* gene, resulting in a VitC dietary dependency similarly as in humans(*20*). Earlier studies utilizing this model showed embryonic lethality at late gestation due to brain and cardiac defects after more than 20 days of VitC deprivation (*21, 22*). Recently, a series of high-profile studies revealed new critical roles of VitC in hematopoietic stem cell function, suppressing leukemia progression, and supporting fetal germ cell development through the regulation of TET activity (*23–25*). However, in these studies using the C57Bl/6J (B6) *Gulo*^tm1Mae^ strain KOs, maternal VitC deficiency was compatible with grossly normal embryonic development until E15.5(*23, 24*).

During early post-implantation embryonic development in the mouse, *Tet1* is the only TET dioxygenase gene with detectable expression in the epiblast (*26*). In our *Tet1*^tm1Koh^ mutant mouse model, complete *Tet1* KO results in cranial NTDs at strikingly much higher penetrance (>60%) in non-inbred than in inbred strains (25%) such as B6 congenics, suggesting a strong impact of genetic backgrounds on the phenotypic expression (*27*). Given that VitC is a co-factor for TET1 activity, we ask whether strain genetic background is also a major confounder of the impact of VitC deficiency on embryonic development, *i.e.* whether the absence of early peri-gastrulation defects in VitC-deficient B6-*Gulo* KO mice would manifest more severe phenotypes in alternative strains. By doing so, we demonstrate dose-dependent effects of VitC deficiency on a spectrum of fetal pathologies ranging from gastrulation defects to NTDs in outbred stocks and incipient congenic strains, in association with

DNA hypermethylation signatures. Further, we define the vulnerable developmental windows during which sufficient VitC status is critical to prevent birth defects as well as longer-term health issues.

## Results

### Maternal VitC deficiency results in strain-dependent malformations and developmental delays in embryos

We have previously created *Tet1* mutant mouse strains that produce high prevalence of NTDs in fetuses by outcrossing B6-*Tet1*^tm1Koh^ to CD1 stocks or backcrossing to a 129 substrain (129S6, also known as 129/SvEv) for 6 to 10 generations (*27*). Using NTD as a trait, we have further performed strain intercrosses between 129S6 and B6 genetic backgrounds for genotype-phenotype correlation analysis in *Tet1* KO embryos, identifying a highly significant quantitative trait locus for NTD risk on chromosome 9 in the 129S6 genome (*28*). To determine if similar strain-dependent disease susceptibilities are observed in vitamin C deficiency, *Gulo*^tm1Mae^ mice were also outcrossed to CD1 (>3 generations), or backcrossed to the 129S6 background for 6-7 generations to create a new 129S6.B6 incipient congenic strain. On the three distinct genetic backgrounds (B6, CD1 and 129S6.B6), we scored phenotypes of embryos at specific developmental stages post-neural tube closure (NTC) (E10.5-E11.5), during NTC (E8.5-E10.5) and during gastrulation (E6.5-E8.5) following withdrawal of VitC from *Gulo*^-/-^ dams three days prior to timed mating and throughout pregnancy (Figure 1a). By performing high performance liquid chromatography-tandem coupled mass spectrometry (HPLC-MS/MS) to detect L-ascorbate levels in the plasma of female mice during the time-course of VitC withdrawal, we confirmed that plasma ascorbate levels have plummeted to background levels by 10 days of VitC withdrawal, which corresponds to a E3.5-E6.5 developmental window in our experimental design, in B6 and 129S6.B6 strains (Supplementary Figure 1a).

**Figure 1.**
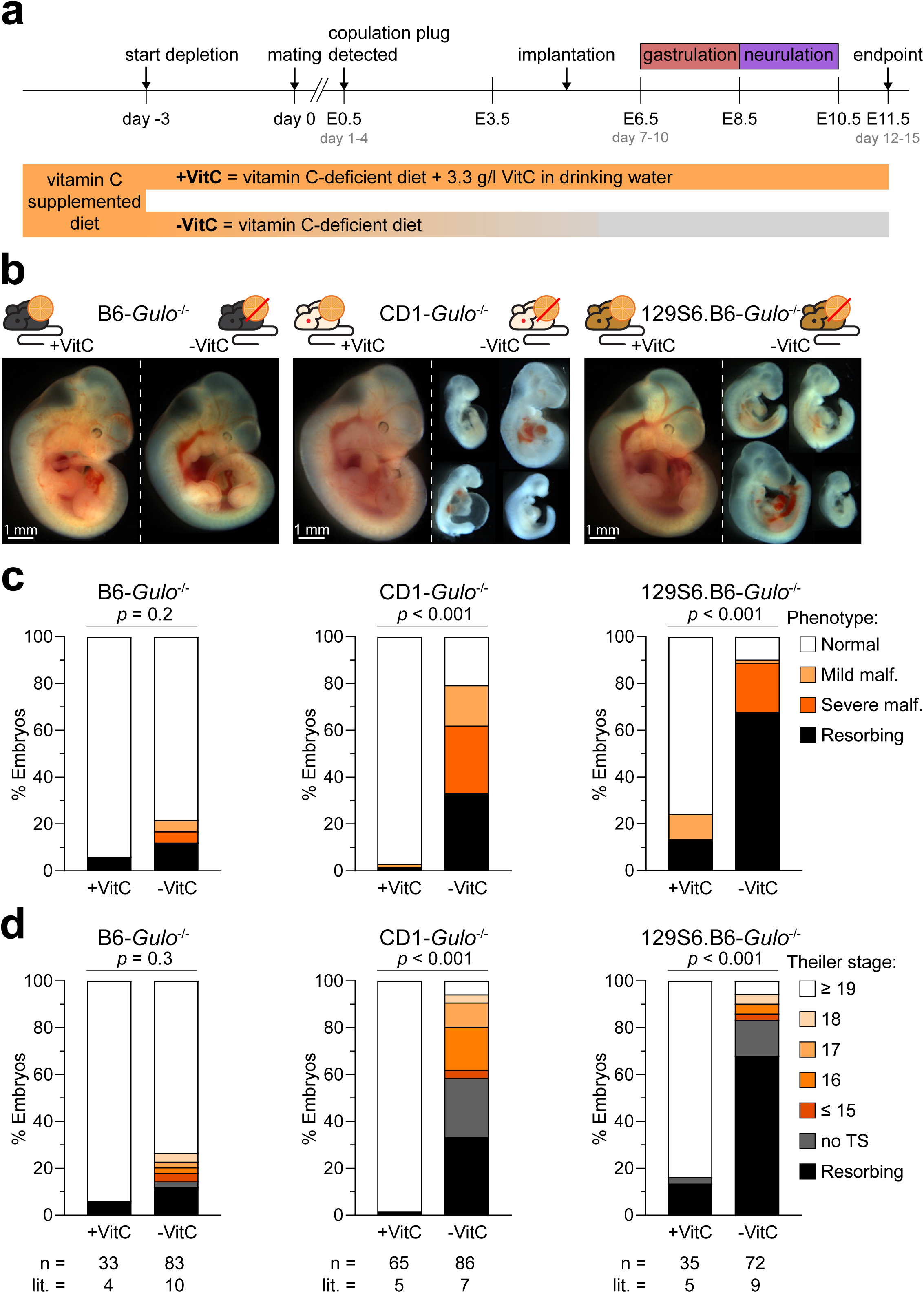
Maternal VitC deficiency results in strain-dependent malformations and developmental delays in embryos. **a,** Experimental schematic for VitC withdrawal (-VitC) and supplementation (+VitC). *Gulo*^-/-^ mice are maintained on diet containing 1% stabilized VitC. Three days prior to timed mating, diet was changed to regular chow (without VitC). +VitC females are supplemented with 3.3 g/l VitC in the drinking water. **b,** Images of representative +VitC and -VitC *Gulo*^-/-^ embryos on inbred B6 (left), outbred CD1 (center), incipient congenic 129S6.B6 (right) genetic backgrounds collected at E11.5. Overview per strain is assembled from individual embryo images, with dashed line separating embryos from different treatments. Scale bar indicates 1 mm. **c, d,** Phenotype (c) and staging (d) scores of +VitC and -VitC E11.5 *Gulo*^-/-^ on the indicated backgrounds. “mild malformation” (malf.) = one gross structural malformation; “severe malformation” = 2 or more types of malformations or grossly malformed; “Resorbing” = empty decidua or embryos in the process of being resorbed; “no TS” = not possible to assign a Theiler stage. n = number of embryos per condition; litters (lit.) = number of litters per condition. Pairwise comparisons are done using Chi-square test.

As reported previously, B6-*Gulo*^-/-^ mouse embryos subjected to maternal VitC withdrawal can develop until E13.5 without gross abnormalities (*21, 23*). Based on severity of embryonic defects and developmental Theiler stages (TS), less than 20% of VitC-deprived (-VitC) B6-*Gulo*^-/-^ embryos appeared malformed or were resorbed in our experiments (terminated at the endpoint of E11.5) (Figure 1b, c) and the majority scored as TS ≥ 19, comparable to VitC-supplemented (+VitC) embryos (Figure 1d). In stark contrast, on both non-inbred CD1 and129S6.B6 backgrounds, maternal VitC deficiency resulted in highly penetrant embryo deformities and developmental delays by E11.5 (Figure 1b and Supplementary Figure 1b). We observed various malformations: craniofacial malformations, including morphological distortion at the midbrain-hindbrain juncture (“hindbrain abnormality’), malformed and hypertrophic heart, grossly reduced body size, and resorbing embryos (Supplementary figure 1c, see Methods for scoring descriptions): > 80% of all CD1 and nearly 90% of 129S6.B6 VitC-deprived embryos were scored as malformed or in the process of resorption; > 90% scored TS < 19 (Supplementary Figure 1d), and > 50% were so grossly malformed that a TS assignment was not possible (Figure 1c,d). In the most severely affected cases, decidua were empty or only contained resorbing embryos, and placental tissues were highly hemorrhagic, suggesting that some embryo defects may be secondary to placental dysfunction (Supplementary figure 1e). These phenotypes are reproducible across random matings and were almost fully rescued when dams were supplemented with 3.3 g/L VitC in drinking water throughout pregnancy (Fig 1b-d and Supplementary Figure 1a). In our experimental design, the duration between mating and detection of copulation plug in dams varied from 1-3 days (corresponding to 4-7 days of VitC deprivation at the time of fertilization). We did not observe any clear trend in correlation between duration of VitC deprivation and phenotype severity, although in two CD1 litters that experienced the longest period of VitC withdrawal (7 days before plug detection), over 70% of embryos were dead or dying at the endpoint (Supplementary Figure 1f). We observed a significant increase in the numbers of resorbed embryos per litter and decreased litter size in CD1 and 129S6.B6, but not in B6 litters (Supplementary Figure 1g, h). Collectively, these results confirm that mice that are genetically heterogeneous (CD1) or harbor unknown modifier risk loci for congenital defects (129S6.B6) are highly sensitive to perturbation by VitC deficiency during embryonic development and that VitC supplementation can overcome embryonic lethality. To assess the impact of VitC withdrawal on non-chromatin (eg. collagen) hydroxylation, we performed histological Sirius red staining of embryo sections for collagen and ruled out disruption of collagen hydroxylation within 2 weeks of VitC withdrawal (Supplementary Figure 2a), in agreement with a previously published report that collagen synthesis in mice is largely independent of VitC (*29*) To characterize the time-course of phenotypic expression in VitC-deprived *Gulo*^-/-^ mice, we collected embryos at earlier developmental timepoints. At E8.5 VitC-deprived embryos appeared mostly normal in all three backgrounds, with only 5-10% found to be resorbing in CD1 and 129S6.B6 litters. However, > 50% of VitC-deprived CD1 embryos had fewer than 3 somites, compared to only 20% in VitC-supplemented (+VitC) litters, suggesting already a developmental delay that reached statistical significance (Supplementary Figure 2b). By E9.5, > 90% of VitC-deprived CD1 embryos staged at TS 14 or below and were considered developmentally delayed compared to normal TS 15-16 at E9.5, or were in the initial stages of being resorbed, while only < 10% of VitC+ embryos were delayed (Supplementary Figure 2c). These observations suggest that the gross developmental phenotypes caused by VitC deprivation in outbred mice already manifest by post-gastrulation.

### Genetic differences result in distinct differential DNA methylation patterns in embryonic headfold tissues upon VitC withdrawal

Having determined the congenic B6- and incipient congenic 129S6.B6-*Gulo* KO strains as displaying low (resistant) versus high (sensitive) penetrance of embryonic defects, respectively, in response to VitC withdrawal, we collected E8.5 headfold tissues from VitC-supplemented (+VitC) and VitC- deprived (-VitC) embryos from both strains for DNA methylome profiling by whole genome bisulfite sequencing (WGBS) (Figure 2a and Supplementary Figure 3a). To account for litter effects as a confounder, we selected no more than one male and one female embryo per litter as biological replicates in our analysis, with an exception in the last litter collected in the 129S6.B6 VitC-deprived group when no stage-matched females were available. Embryos were matched by somite counts of 4-6 to detect molecular changes preceding the appearance of gross morphological defects, although in two litters from VitC-deprived 129S6.B6 mice, embryos appeared to be in an early stage of resorption. In all, we selected headfold tissues from 4 embryos (male and female) in each of 4 treatment groups: +VitC versus -VitC treatments in two strains (B6 and 129S6.B6), stage-matched for normal morphology across groups, and added two more samples from the 129S6.B6 VitC-deprived litters displaying resorbing phenotypes (labelled -VitC pheno) (Supplementary Figure 3b).

**Figure 2.**
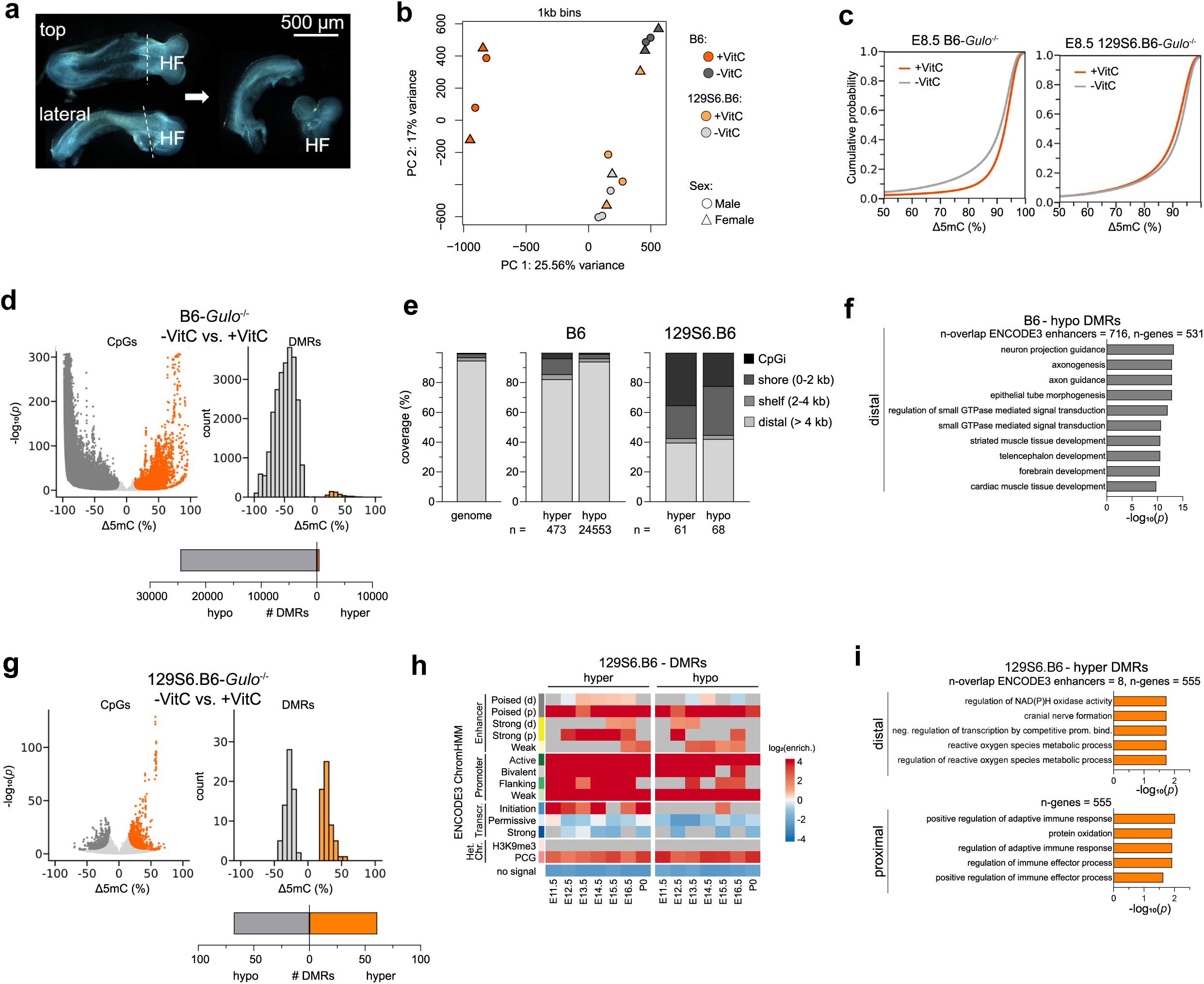
Genetic differences result in distinct differential DNA methylation patterns in embryonic headfold tissues upon VitC withdrawal. **a**, Representative E8.5 embryo used for dissection in top and lateral view (left), and the dissected headfold (right). Dissection plane is indicated with dotted line. HF, headfold. Scalebar indicates 500 µM. **b**, **c**, PCA (b) and cumulative distribution (c) plots of CpG methylomes from E8.5 headfolds, collected from +VitC and -VitC B6- and 129S6.B6-*Gulo*^-/-^ embryos. Both plots were calculated using average CpG methylation levels in 1-kb bins and a minimal coverage of 10. **d**, Volcano plot of differentially methylated CpGs, orange and grey highlight indicating those with significant gain or loss, respectively, in -VitC relative to +VitC, >10% and FDR < 0.001 (left), histogram of DMR distribution by differences in CpG methylation levels in intervals of 10% (right), and number of hyper and hypo DMRs (bottom) in B6-*Gulo*^-/-^ headfold tissues. **e**, Distribution of hyper and hypoDMRs by proximity to CpG islands (CpGi) in both B6- and 129S6.B6-*Gulo*^-/-^ headfolds. Genome reference distribution of genomic features are shown on the left. **f**, Top 10 enriched GO terms of B6 hypoDMRs that overlap with ENCODE3 enhancers. **g**, Plots of significantly differentially methylated CpGs, distribution and numbers of DMRs in 129S6.B6-*Gulo*^-/-^ headfolds, as in (d). **h**, log2-fold enrichment of 129S6.B6 DMRs at functional genomic elements active during mouse fetal development from E10.5 to P0 as classified by 15 chromatin states in ENCODE3. **i**, Top 5 enriched GO terms of 129S6.B6 hyper DMRs that overlap proximally and distally with genes associated with ENCODE3 enhancers.

WGBS analysis showed the latter two “-VitC pheno” samples had lower coverage, higher duplication rates and lower global global CpG methylation levels compared to all other samples (Supplementary Figure 3c, d), so that they were considered separately from the 4 morphologically “normal” samples in the 129S6.B6 VitC-deprived group in downstream differential analysis. Principal component analysis (PCA) of CpG methylation patterns showed clearly distinct DNA methylomes in +VitC headfolds between the two genetic backgrounds (Figure 2b). VitC withdrawal resulted in dramatic shifts in all B6 methylomes along the first principal component (PC1), in contrast to a more subtle shift in the 129S6.B6 methylomes of “normal” headfolds in a different direction along PC2. The two 129S6.B6 -VitC pheno samples were outliers when included in the PCA (Supplementary Figure 3e), further justifying their exclusion in the analysis based on normal morphology. Cumulative probability distribution curves of CpG methylation levels revealed a pronounced global trend of DNA hypomethylation in VitC-depleted B6 embryos, which are refractory to early defects, but a slight shift towards hypermethylation in VitC-deprived 129S6.B6 embryos, which do subsequently succumb to malformation (Figure 2c).

In VitC-deprived B6-*Gulo*^-/-^ tissues, a reduction in global CpG methylation levels relative to VitC controls were detectable across CpG islands (CpGi) shelves and shores, gene bodies and repetitive sequences (Supplementary Fig. 3f). 482,184 CpGs were differentially methylated (467,533 gain and 14,651 loss), corresponding to 25,026 differentially methylated regions (DMRs) of which 98.1% (24,553) showed collective loss of DNA methylation (Figure 2d). These prevalent “hypo-DMRs” in the B6 strain are enriched for distal enhancers regulating developmental genes as defined by ENCODE3 (*30*), of which the top gene ontology (GO) terms are in “neuron projection guidance” and “axonogenesis” (Figure 2e,f). These may affect postnatal rather than pre-natal development. In contrast, 129S6.B6 -VitC embryos displayed more subdued differences from +VitC controls, with similar global CpG methylation levels across genomic features (Supplementary Figure 3g). Nonetheless, the VitC-deprived samples that appeared morphologically normal like +VitC controls displayed 2,732 significantly differentially methylated CpGs, 1,565 with gain and 1,167 with loss, which spanned 61 hyperDMRs and 68 hypoDMRs respectively (Figure 2g). Both hypo- and hyper-DMRs in the 129S6.B6 strain are strongly enriched at CpG islands (CpGi) and shores (Figure 2e), particularly at functional regulatory elements ( promoters and enhancers) which are active throughout mouse fetal development from E11.5 to P0 as annotated by ENCODE3 (*31*) (Figure 2h and Supplementary Figure 3h). Whereas there was no significant association of 129S6.B6 hypoDMRs with GO terms, hyperDMRs overlapped proximally with 555 genes associated with “regulation of adaptive immune response” and distally with 8 ENCODE3 enhancers associated with GO terms including “reactive oxygen species metabolic process” and “cranial nerve formation” (Figure 2i). Given their low numbers, these 129S6.B6-specific hyperDMRs enriched poorly for specific developmental pathways, but they nonetheless overlapped significantly with ENCODE3 enhancers linked to 2 genes *Tfap2a* and *Cyba*. Interestingly, *Tfap2a* encodes an AP-2 alpha transcription factor involved in cell division, apoptosis, and neural tube closure during embryonic development(*32*), suggesting that DNA hypermethylation at this gene may be linked with NTD phenotypes later. The two “-VitC pheno” samples showed more extensive differential methylation trending towards hypermethylation compared to VitC+ controls, resulting in 387 hyperDMRs, but these were predominantly at distal sites (Supplementary Figure 3i, j).

To relate the DNA methylation changes with gene expression, we collected E8.5 headfold tissue RNA from additional embryos (B6,+VitC: n=5, 3 males and 3 females from 3 litters; B6,-VitC: n=8, 5 males and 3 females from 3 litters; 129S6.B6,+VitC, n=8, 5 males and 3 females from 4 litters;129S6.B6,-VitC: n=7, 4 males and 3 females from 4 litters; Supplementary Figure 4a) for low-input RNA-sequencing. Based on PCA, one outlier sample (B6, +VitC, female) was excluded from downstream analysis (Supplementary Figure 4b). All other samples separated clearly by strain along the first principal component (PC1), whereas +VitC and -VitC samples per strain, especially 129S6.B6, still overlapped along PC2 (Supplementary Figure 4c). Differential expression analysis by - VitC versus +VitC treatment per strain identified 253 differentially expressed genes (DEGs) in B6, 119 up- and 134 down-regulated in roughly equal proportion, and only 62 in 129S6.B6, of which most (48) were down-regulated (Supplementary Figure 4d). Both up- and down-regulated DEGs in VitC- deprived B6 headfolds preferentially resided near DMRs and were significantly associated with developmental GO terms including “pattern specification process” and “regionalization” (Supplementary Figure 4f). These results are paradoxical because disruption of TET dioxygenase activities by VitC withdrawal is expected to drive DNA hypermethylation and gene silencing. Instead, we observed a global hypomethylation in the B6 background that may affect expression of pattern specification genes, but nonetheless did not result in detectable embryonic phenotypes until E11.5. In contrast, DEGs in VitC-deprived 129S6.B6 headfolds were not preferentially within 100 kb of DMRs. Those down-regulated were associated with GO terms in meiosis, in line with a primary impact maternal VitC deficiency on germ cell development(*23*), but not with gross developmental terms (Supplementary Figure 4g). For instance, hyperDMRs at *Tfap2a* and *Cyba* were not associated with any significant change in gene expression. Collectively, these results show that the genetic background is a major determinant of the epigenome response to VitC withdrawal, with a hypermethylation signature strikingly associated with strain susceptibility to congenital malformations. HyperDMRs in the susceptible 129S6.B6 strain are enriched at functional gene regulatory elements previously validated during fetal organogenesis post-gastrulation (E8.5) but yet are largely latent marks that do not directly affect gene expression.

### Sub-optimal levels of VitC can trigger a spectrum of embryonic pathologies associated with DNA hypermethylation

Using CD1-*Gulo* KO mice, we performed dose-titration of VitC supplementation to assess the effects on embryo phenotypes at the endpoint of E11.5 (Figure 3a). In agreement with previous studies, 330 mg/l is the minimal dose required to sustain normal development in over 90% of offspring(*33*). Reduction to 33 mg/L caused partially penetrant embryonic malformations, including resorptions, to reach 40% (Figure 3b, c). Interestingly, the intermediate 100 mg/l VitC dose resulted in malformation in about 15% of stageable embryos, including NTDs specific to the cranial regions as a new phenotype (Figure 3c). HPLC-MS/MS measurements of plasma ascorbate levels in pregnant dams collected at the experimental endpoint in these VitC dose-response treatments confirmed that the 330 mg/l dose resulted in plasma ascorbate levels equivalent to that in WT mice (5 μM). Detectable plasma ascorbate levels dropped by half in the 100 mg/l dose treatment and to baseline by the 33 mg/l dose (Figure 3d). These results provocatively suggest that a moderate reduction in VitC status (at the threshold below 5 μM in circulation) can trigger a spectrum of fetal pathologies.

**Figure 3.**
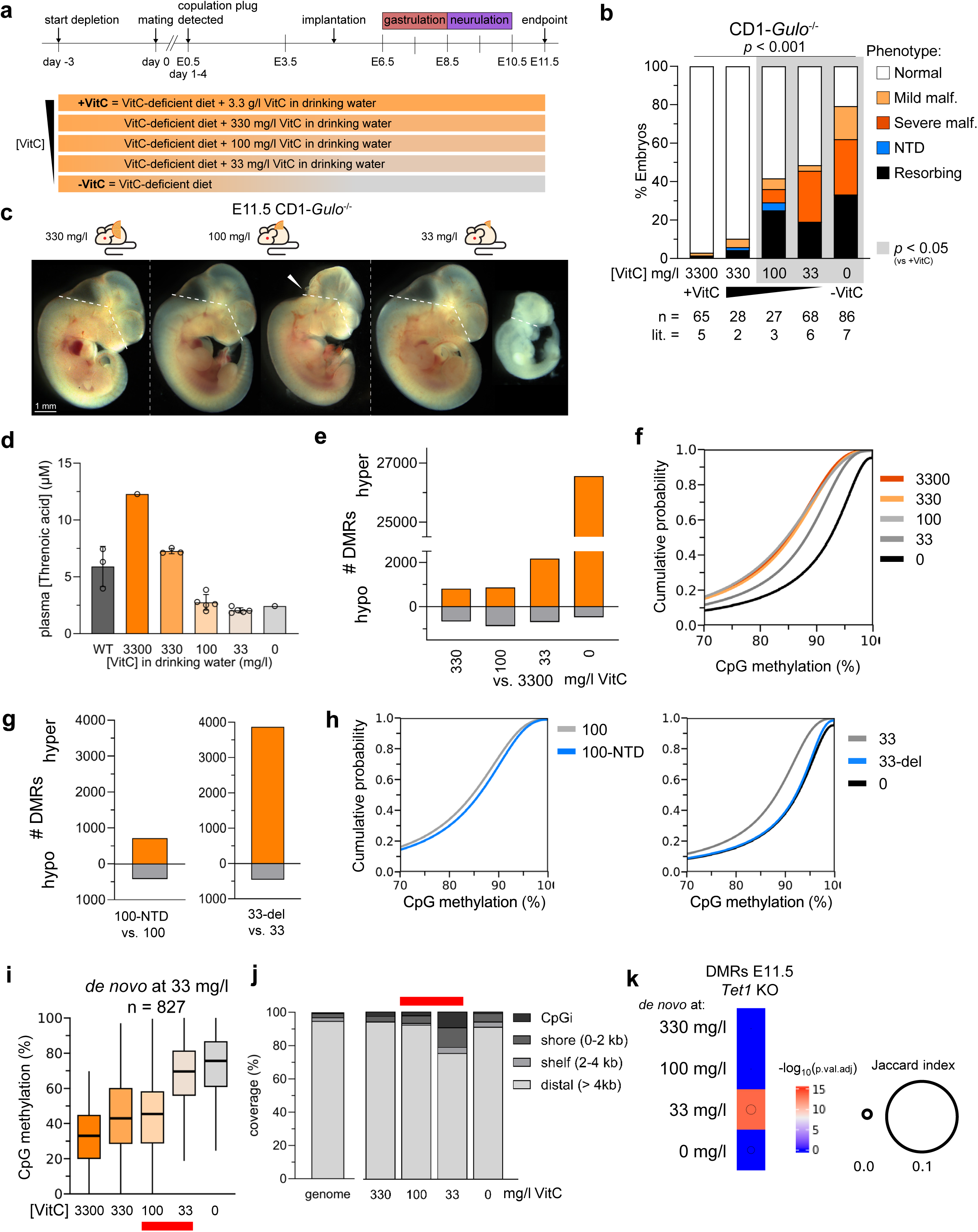
Suboptimal levels of VitC can trigger a spectrum of embryonic pathologies associated with DNA hypermethylation**. a**, Experimental scheme for VitC dose-titration using CD1-*Gulo*^-/-^ mice. **b**, Embryo phenotype scoring. Malf., malformed. Overall *p* from Chi Square test is shown above. Pairwise comparison per dose treatment with +VitC (3300 mg/l) control is adjusted with Bonferroni multiple testing correction. Grey shades indicate dose-range at which there was a significant increase (Chi Square test *p* < 0.05) in malformations that were detected compared to +VitC control. **c**, Representative E11.5 embryos used in WGBS. Arrowhead marks NTD. Dashed white lines indicate planes of dissection to collect brains. We collected the entire brain, consisting of the hindbrain (HB), midbrain (MB), and forebrain (FB). **d**, Plasma level of total ascorbate detected as the degradation product threnoic acid by LC-MS/MS. **e**, Number of hypo (grey) and hyper (orange) DMRs in E11.5 brains (all with normal morphology until the lowest dose of 33 mg/l) from pairwise comparison of each treatment with +VitC control. **f**, Cumulative probability distribution of mean CpG methylation levels in 1-kb bins with a minimal coverage of 10 in E11.5 brains at each indicated VitC dose. **g**, Numbers of hypo and hyper DMR in E11.5 brains from comparison between littermate embryo pairs with phenotype (NTD, or del., delayed) versus normal. **h**, Cumulative distribution plots (as in f) for CpG methylation levels in E11.5 brains from littermate embryo pairs with and without phenotypes. **i**, Dose-response of CpG methylation levels in a subset of *de novo* hyperDMRs at 33 mg/l VitC, defined as regions exhibiting a significant increase in CpG methylation specifically upon a half-logarithmic VitC dose reduction from 100 to 33 mg/l (red bar) during the graded dose reduction. The box represents the interquartile range (IQR) and the line within indicates the median. **j**, Distribution of *de novo* hyperDMRs per VitC dose by CpG island proximity. Red bar marks “sensitive” dose range for elevated penetrance of severe malformations. **k**, Jaccard overlap of *de novo* hyperDMRs per VitC dose with hyperDMRs in E11.5 *Tet1* KO brain(*27*).

To examine the DNA methylome, we selected E11.5 embryos staged at TS19-20 with normal morphology from each VitC dose exposure from 3300-33 mg/l and dissected the brains for WGBS (Figure 3c). A littermate embryo with NTD at 100 mg/l VitC and two growth delayed embryos with intact brains at 33 and 0 mg/l VitC were also included (Supplementary Figure 5a). PCA of CpG methylation profiles revealed global methylome shifts along both PC1 and PC2 at doses 100 mg/l and below, and further shifts in one direction on PC1 in embryos with phenotypes (Supplementary Figure 5b). Compared to the E11.5 brain supplemented with high dose VitC at 3.3 g/ml (reference control), moderate VitC dose reductions to 100 mg/l resulted in roughly equal numbers of hypo- and hyper- DMRs in brains with normal morphology (Figure 3e and Supplementary Figure 5c). However, reduction to 33 mg/ml triggered an increase of hyper DMRs by over 2-fold, even in an embryo that appeared normal. Notably, hypoDMR numbers remained relatively constant in response to VitC dose reduction, but hyperDMRs far exceeded hypoDMRs in numbers once VitC dose was lowered to 33 mg/l. E11.5 brains exposed to 33 mg/l VitC and below, even when morphologically normal, showed evident global hypermethylation and an increase in CpG methylation levels across CpG island (CpGi) flanking regions, gene bodies and retrotransposon repeats (Figure 3f and Supplementary Figure 5d,e).

In the brains from embryos with phenotypes, the numbers of hyper DMRs increased further, suggesting that hyper DMRs accumulation correlates with the severity of embryonic defects (Figure 3g,h and Supplementary Figure 5c). These phenotype-associated hyperDMRs were enriched across distal intragenic regions, but depleted from CpGi shores (Supplementary Figure 5d, f, g). In the NTD-affected brain compared to normal brain at 100 mg/l VitC exposure, hyperDMRs enriched for ENCODE3 fetal enhancers associated with genes described by GO terms in “cell-substrate adhesion” and “neuron death” (Supplementary Figure 5h, i). In the brains from delayed versus normal embryos at 33 g/l VitC exposure, hyperDMRs overlapped even more extensively with enhancers associated with GO terms in forebrain development and axonogenesis (Supplementary Figure 5j, k). The GO terms enriched by the hyperDMR signatures concur with the phenotypes observed.

RNA-seq of E11.5 brains from littermate embryos (in duplicates per dose and phenotype; Supplementary Figure 6a) showed a segregation in the transcriptomes between embryos exposed to VitC doses either higher or lower than 100 mg/l, in line with detection of >100 DEGs in normal brains when VitC dose dropped to 100 mg/l (Supplementary Figure 5b) and a significant increase in the incidence of embryonic phenotypes (Figure 3b). DEGs appeared in almost equal ratios of up- and down-regulation, increased by 2-fold with VitC dose reduction to 33 mg/l in brains of normal morphology, and a further 5-fold in the brains from embryos with gross developmental delays (Supplementary Figure 6c,d). In the embryos exposed to 100 mg/l VitC, appearance of NTDs resulted in 16 DEGs, of which 13 were down-regulated (Supplementary Figure 6d). In every condition, DMRs generally do not overlap with DEGs. While this could be because we sampled brains from different individuals in the two read-outs, the threshold for globally detectable changes in dose-response to VitC reduction is clearly different for the transcriptome versus the DNA methylome even in stage-matched embryonic tissues with normal morphology – 100 ng/ml for widespread DEGs to be detectable and 33 mg/ml for DNA hypermethylation, suggesting that VitC status may affect gene expression and DNA methylation by distinct mechanisms.

We further defined subsets of hyperDMRs that accrue *de novo* with each step of VitC dose reduction and stayed hypermethylated with subsequently lower dose(s), calling these *de novo* DMRs corresponding to each dose (Supplementary Figure 7a, b and Figure 3i). Dose reductions down to 100 mg/l generated DMRs that did not enrich significantly for any GO terms, suggesting that these are mostly stochastic DMRs at distal loci. Interestingly, *de novo* DMRs appearing at 33 mg/l VitC, the dose triggering an increased rate of severe malformations in stageable embryos (Figure 3b), were enriched for CpGi and shores, and also at ENCODE3 annotated fetal promoters and enhancers compared to the other classes of *de novo* DMRs (Figure 3j and Supplementary Figure 7c). They also overlapped significantly with hyper DMRs in E11.5 *Tet1* KO brains, suggesting that they occur from TET1 loss of function (Figure 3k). GO analysis of these hyper DMRs (*de novo* at 33 mg/l) indicated enrichment for terms in cell development, including “regulation of leukocytes differentiation” and “negative regulation of cellular response to growth factor stimulus” as the top terms. In agreement, the overlap with ENCODE3 enhancers enriched for GO terms in both neuronal and mesenchymal development (Supplementary Figure 7d). These results suggest that even in embryos that appear morphologically normal, a pathologically low VitC status (at 33 mg/l) can leave aberrant DNA hypermethylation signatures at loci regulating all germ layer lineages, coinciding with disruption of TET activity. Upon complete VitC withdrawal (0 mg/l), *de novo* hyperDMRs extended to enhancers associated with “axogenesis” and “forebrain development”, in line with the overt embryonic defects afflicting almost all embryos (Supplementary Figure 7e).

### Critical window of sensitivity to VitC deficiency coincides with gastrulation

To define the developmental time-window of sensitivity to VitC depletion, we performed a series of timed VitC rescue during peri-gastrulation stages, by injecting pregnant CD1-*Gulo*^-/-^ dams with a single dose of 4 g/kg sodium ascorbate in PBS, between E4.5-E9.5. This high single dose injection was previously shown to be well tolerated by mice (*25*). After this acute rescue treatment, pregnant dams continue to receive supplementation with 3.3 g/L VitC in the drinking water until the experimental endpoint of E11.5 (Figure 4a). VitC rescue was most effective when initiated at E4.5- E7.5, which reduced developmental delays and malformations to < 25%, but only partially effective by E8.5 (> 70% delays, malformations and resorbing embryos), and ineffective after E9.5 (> 95% penetrance) (Figure 4b). These results suggest that the sensitive window for disruption of dioxygenase activity to cause birth defects occurs during gastrulation, after which full VitC supplementation is no longer effective. These findings are completely consistent with a previously published study that has shown gastrulation defects in genetic ablation of several epigenetic regulators (*12*). Interestingly, we observed one case of NTD in a CD1-*Gulo*^-/-^ embryos rescued with VitC at E6.5, suggesting that an earlier window of VitC deprivation pre-gastrulation may contribute to some NTDs (Figure 4c).

**Figure 4.**
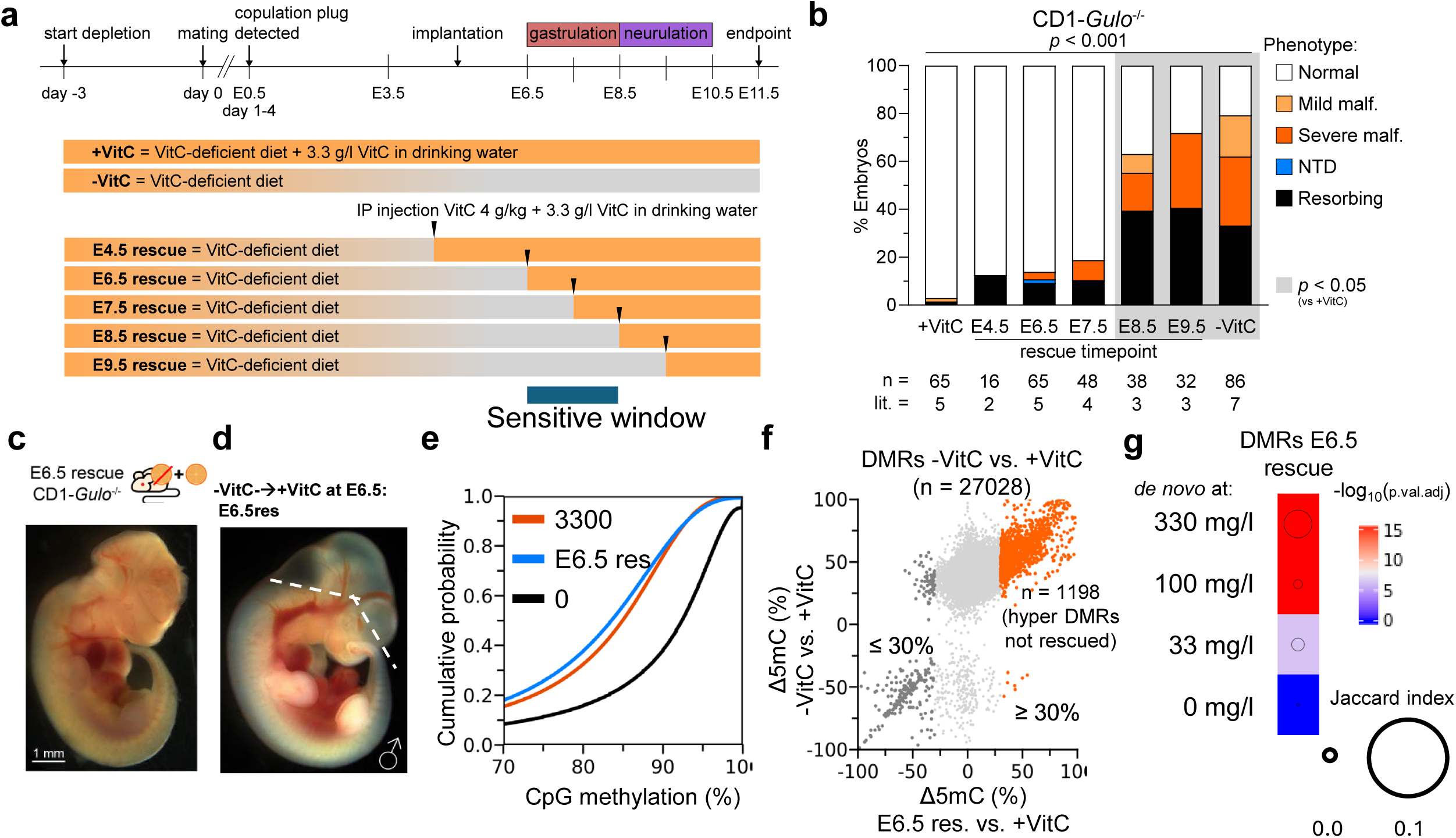
Critical window of sensitivity to VitC deficiency coincides with gastrulation. **a**, Experimental schematic for VitC re-supplementation time-course. In each rescue experiment, VitC- deprived pregnant dams were injected intraperitoneally with 4g/kg VitC at the indicated timepoint and subsequently sustained with 3.3 g/l VitC via the drinking water. **b**, Penetrance of phenotypes in E11.5 -VitC CD1-*Gulo*^-/-^ embryos rescued with VitC at the indicated timepoints. Overall *p* from Chi Square test is shown above. Grey shades indicate timed-rescue treatments in which significant increase in malformations were detected (not effectively rescued) in pairwise comparison to +VitC control (Chi square p < 0.05 adjusted with Bonferroni multiple testing correction). **c**, A VitC-deficient CD1-*Gulo*^-/-^ embryo rescued by high-dose VitC re-supplementation at E6.5, with an NTD. **d**, A fully rescued “normal” CD1- *Gulo*^-/-^ embryo after VitC re-supplementation at E6.5. Dashed line indicates dissection plane above which brain tissues were collected for WGBS. **e**, Cumulative distribution of mean CpG methylation levels in 1-kb bins with a minimal coverage of 10, in E11.5 brains exposed to 3.3 g/l (+VitC), 0 mg/l (-VitC) and re-supplementation at E6.5 (E6.5 res). **f**, Scatter plot of CpG methylation changes at DMRs in an E11.5 brain caused by complete VitC withdrawal (-VitC) compared to changes after rescue at E6.5, relative to +VitC control. Hyper DMRs resistant to VitC rescue are marked in orange in the top right quadrant. **g**, Jaccard overlap of resistant hyperDMRs with *de novo* hyperDMRs per VitC dose.

DNA methylome analysis of an E11.5 brain from an embryo rescued by VitC re-supplementation at E6.5 (E6.5 res) showed a nearly complete restoration of DNA methylation to that of a fully VitC- supplemented control, in line with the normal morphology of the rescued embryo (Figure 4d,e). When compared to fully VitC-supplemented E11.5 brain, the E6.5 rescued brain showed a slight trend of global DNA hypomethylation, with global CpG methylation levels slightly reduced (Figure 4e and Supplementary Figure 5d). DMR analysis revealed 1713 hypo-DMRs, indicating regions overly demethylated relative to +VitC control, which slightly out-numbered 1257 hyper-DMRs indicating regions not fully demethylated (Supplementary Figure 8a). These were all depleted of CpGi and shores, with an enrichment of hyperDMRs for distal intergenic regions (Supplementary Figure 8b,c). Against the set of all DMRs between fully supplemented (+VitC) and complete withdrawal (-VitC) treatment, a majority (94.6%) were normalized by VitC re-supplementation except for 1198 hyper DMRs which resisted VitC rescue (Figure 4f). These “resistant” hyperDMRs (i.e. not rescued) overlapped significantly with *de novo* hyper DMRs induced at 330 mg/l and 100 mg/l VitC (Figure 4g), suggesting that they are loci highly sensitive to mild VitC insufficiency. The top GO term enriched was “sodium-independent organic anion transport”, followed by several terms related to “acid transport” and “immune response” among the top 10 (Supplementary Figure 8d). RNA-seq analysis of VitC-deprived brains rescued at E6.5 compared to VitC+ controls identified 20 up- regulated and 32 down-regulated DEGs (Supplementary Figure 8e). Top GO terms enriched by down- regulated genes are all related to “antigen processing and presentation” (Supplementary Figure 8f), suggesting that these resistant hyperDMRs are repressing gene pathways associated with the immune response already at E11.5. These results imply that an earlier window vulnerable to insufficient VitC could affect offspring health and immunity despite averting birth defects.

### VitC deficiency induces TET1-dependent and independent epigenetic changes

In both CD1 and 129S6.B6 mice, the phenotypes of the VitC-deprived *Gulo*^-/-^ embryos were significantly more severe than observed in *Tet1*^-/-^ embryos. In the latter, the phenotype consists of an NTD and lacks associated gross structural malformations and severe growth restrictions. These observations are in line with our expectation that VitC deficiency may disrupt additional dioxygenases, including JMJC domain-containing histone demethylases, during gastrulation. We asked to what extent the phenotypes observed in the sensitive backgrounds in VitC-deprived litters are attributable to the disruption of TET1 function. To answer this question, we crossed CD1-*Gulo*^tm1Mae^ to the CD1-*Tet1*^tm1Koh^ strain (a double mutant strain henceforth referred as TG) and evaluated how developmental phenotypes resulting from VitC withdrawal are dependent on *Tet1* genotypes in a *Gulo* KO background (*Tet1*^tm1^;*Gulo* KO)(*27*) (Figure 5a).

**Figure 5.**
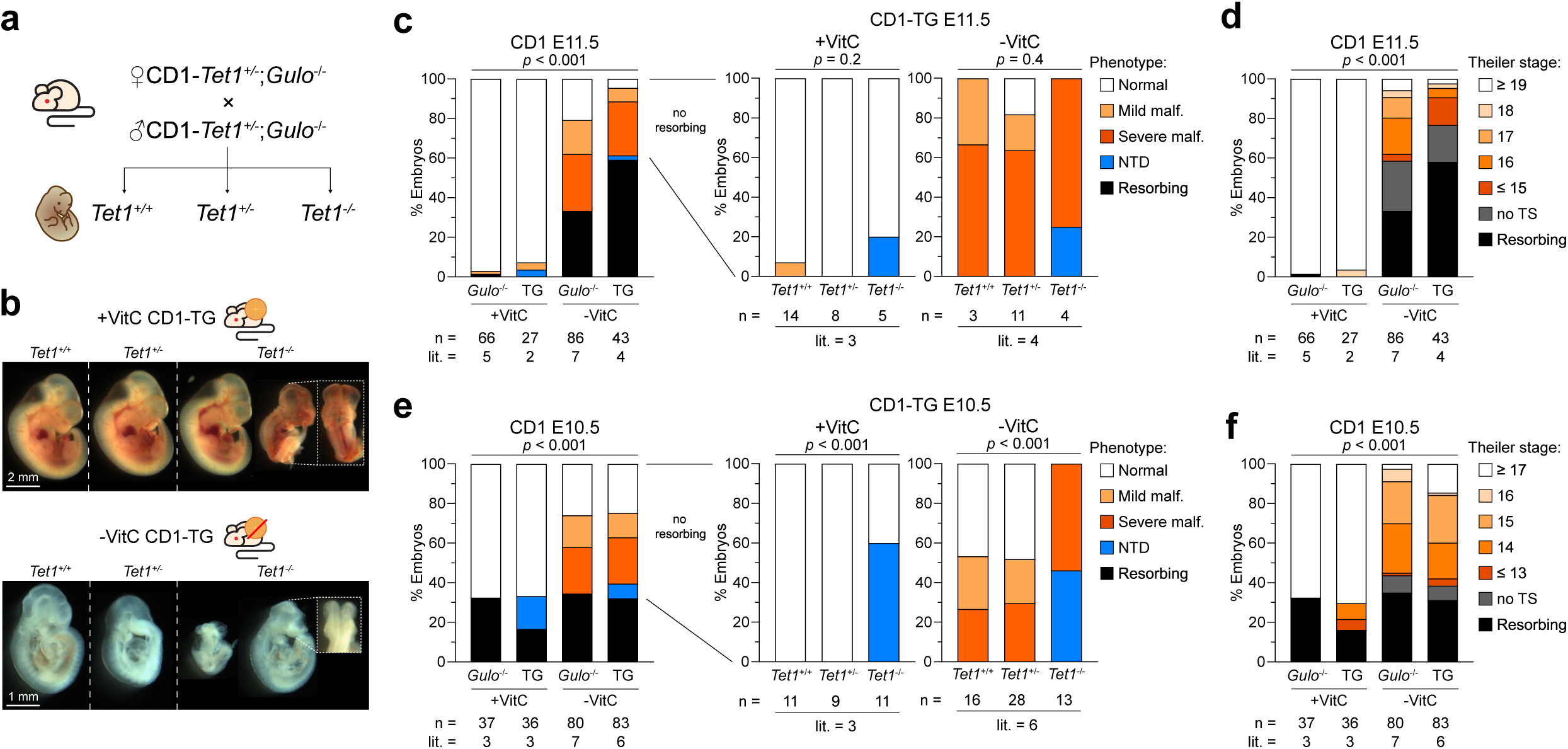
Maternal *Tet1* haploinsufficiency exacerbates -VitC deficiency phenotype. **a**, Experimental timed-mating scheme to generate CD1-*Gulo*^-/-^ embryos with different *Tet1* genotypes. **b**, Representative images of E11.5 +VitC and -VitC CD1-*Gulo*^-/-^;*Tet1*^tm1Koh^ (TG) embryos classified by *Tet1* genotypes. Overview panel per treatment/litter is assembled from individual embryo images, with dashed line separating embryos of different *Tet1* genotypes. The insets show dorsal views of incomplete cranial neuropore closure in *Tet1*^-/-^ embryos in both +VitC and -VitC treatments. **c-f,** Scoring of phenotype penetrance (c,e) and developmental stage (d,f) for +VitC and -VitC embryos from intercrosses between CD1-*Gulo*^-/-^ or CD1- *Gulo*^-/-^;*Tet1*^+/-^ parents at E11.5 (c,d) and E10.5 (e,f). Phenotype scores of CD1-TG embryos in (c,e) are stratified by *Tet1* genotype (right). Statistical comparisons are made using Chi-square test.

Complete VitC withdrawal in TG offspring (comprising *Tet1*^+/+^, *Tet1*^+/-^ and *Tet1*^-/-^ genotypes from both parents which were heterozygous for *Tet1*^tm1^) led to even higher penetrance of malformations and severe developmental delay by E11.5, compared to VitC-depletion in the *Gulo*^-/-^ single-KO (Figure 5b- d). Severe malformations were observed at similar rates in *Tet1*^+/+^, *Tet1*^+/-^ and *Tet1*^-/-^ offspring, suggesting that these phenotypes are independent of the *Tet1* genotype (Figure 5c). Thus, maternal haploinsufficiency of *Tet1*, and not the loss of expression in the offspring, may be exacerbating the phenotypes of offspring due to maternal VitC deficiency. The incidence of resorbing embryos and delayed embryos with a TS ≤ 15 were more frequent in VitC-deprived TG embryos at E11.5 (∼60% resorbed and ∼15% delayed), compared to VitC-deprived *Gulo*^-/-^ (∼30% resorbed and ∼2% delayed) (Figure 5c, d). At E10.5, penetrance of malformed and delayed (TS ≤ 15) phenotypes were still similar in both groups, reaching complete penetrance in phenotypes of severe malformation and NTDs combined only among *Tet1* KO embryos (Figure 5e, f), suggesting that parental *Tet1* haploinsufficiency exacerbates severe malformation, growth arrest and ultimately embryonic lethality in TG embryos post E10.5. Intriguingly, NTDs appeared as a new phenotype in TG embryos only in *Tet1*^-/-^ embryos as expected, but at similar reduced penetrance in both +VitC and -VitC groups (Figure 5c, e). These observations suggest that NTDs specific to *Tet1* KO embryos arise by a different mechanism, such as non-catalytic regulation by TET1, from the most severe gastrulation defects observed in VitC-deprived embryos. Thus, VitC supplementation can rescue all gross structural defects and growth retardation but not NTDs in *Tet1*^-/-^ embryos, consistent with a dominantly non-catalytic function of TET1 during peri-gastrulation development (*11, 34, 35*).

To address whether VitC regulates TET1 catalytic function at mid gestation, we assessed DNA CpG methylation in E11.5 embryonic brain tissues using targeted-bisulfite sequencing, at 3 loci (*Cspg4*, *Gal*, and *Ngb*) we and others have demonstrated to be targets of regulation by the catalytic activity of TET1 (*11*). We previously demonstrated that the promoters of *Cspg4*, *Gal*, and *Ngb* are hypermethylated in neural folds and brains of E8.5 and E11.5 *Tet1*^-/-^ embryos, respectively, prior to gene expression later in development (*11, 36–38*). To determine the extent of DNA hypermethylation at these gene loci caused by VitC withdrawal, we collected E11.5 embryonic brain tissues isolated from VitC-replete (2 litters) and VitC-deprived (3 litters) individual *Gulo*^-/-^ embryos (male and female) of both B6 and 129S6.B6 strains. VitC-deprived B6-*Gulo*^-/-^ embryos were morphologically like +VitC-supplemented B6*-Gulo*^-/-^ embryos (Supplementary Figure 1b). However, many VitC-deprived 129S6.B6-*Gulo*^-/-^ embryos were so malformed that it was not always possible to dissect the brains, limiting us to collecting genomic DNA from the entire embryo in these most severe cases (Supplementary Figure 9c). In both B6 and 129S6.B6 VitC-deprived embryos, significant hypermethylation of the promoters of *Cspg4*, *Gal*, and *Ngb* were detected, compared to VitC-treated embryos (Supplementary Figure 8). Nonetheless, these increases in methylation were much smaller than the increase observed in complete *Tet1*^-/-^ embryonic brains at E11.5 (*11*), suggesting that TET catalytic activity may not be completely abrogated by VitC deficiency. The extent of hypermethylation observed in both -VitC B6-*Gulo*^-/-^ and 129S6.B6-*Gulo*^-/-^ were remarkably similar. Further, there was no difference in methylation between 129S6.B6 brains and whole malformed embryos, or between male and female embryos from both backgrounds (Supplementary Figure 9d). Thus, while VitC withdrawal elevates promoter DNA methylation of these brain-specific genes, the effects are consistent in the two strains and cannot account for the strain-dependent embryonic phenotypes.

To determine whether global histone methylation levels are perturbed by VitC deficiency, we performed Western blot analysis of 5 key histone methylation marks for chromatin states in individual E11.5 brains. These are histone H3 lysine 9 di- and trimethylation (H3K9me2/3), H3K27 trimethylation (H3K27me3), H3K4me3 and H3K4 monomethylation (H3K4me1), all targets of KDMs and important for chromatin biology (*39*). We also assessed H3K27 acetylation (H3K27ac), since it is an important histone mark for active chromatin and act reciprocally to H3K27me3 (*40*). Interestingly, H3K9me2 was the only histone mark showing a global increase in VitC-depleted B6-*Gulo*^-/-^ brains compared to VitC-replete B6-*Gulo*^-/-^ and B6-*Gulo*^+/+^ brains. All other assessed histone PTMs were reduced in VitC-deficient embryonic brains, including both H3K27me3 and H3K27ac. There was no difference between male or female embryos (Supplementary Figure 9e-f). We only analyzed B6-*Gulo*^-/-^ tissues, since there was insufficient material in 129S6.B6-*Gulo*^-/-^ for protein analysis due to the malformations and developmental delays. Nonetheless, these observations suggest that VitC deficiency during embryonic development results in dysregulation of chromatin states in the absence of any morphological abnormalities in B6-*Gulo*^-/-^ embryos.

## Discussion

In this study, we used VitC as a tool to manipulate the JMJC dioxygenase family activities during an early window of mouse embryonic development, creating an experimental model with which to better understand how adverse environmental exposures, including nutritional deficiencies, interact with genetic susceptibilities to alter the epigenome and result in congenital disorders. Using *Gulo*^-/-^ mice bred on non-inbred genetic backgrounds, we show that maternal VitC deficiency results in a spectrum of fetal pathologies ranging from gastrulation defects to NTDs, in a phenocopy of strain-dependent disease penetrance with *Tet1* loss-of-function. These phenotypes were overlooked in previous studies using *Gulo*^-/-^ mice bred onto conventional B6 backgrounds, highlighting that mouse models with higher genetic diversity can provide greater power to understand the impact of gene-environmental interactions on complex disease traits. We show that the intrinsic genetic variation is a strong driver of completely different DNA methylation signatures in response to vitamin C deficiency – widespread DNA hypomethylation by post-gastrulation in the B6 congenic strain, which are refractory to early defects, but a greater proportion of hypermethylation in an alternative incipient congenic 129S6.B6 strain, which do succumb to severe malformation. We established an inverse dose-dependent relationship between maternal VitC status and extent of DNA hypermethylation in outbred embryonic brains even in the absence of morphological phenotypes. DMRs are found at fetal stage gene regulatory elements as early as E8.5 in mice prone to post-gastrulation defects. While these marks appear latent and not affect gene expression, they may contribute causally to developmental defects when coupled to other risk factors. In more severely malformed E11.5 brains from VitC-deprived non-inbred mice, the more widespread prevalence of DNA hypermethylation marks indicates that most of these aberrations are likely the downstream consequences of abnormal development. We determined a pathologically low VitC dose level (33 mg/l) that generates hyperDMRs proximal to fetal developmental genes, which overlap significantly with hyperDMRs in stage-matched *Tet1* KO brains, suggesting that insufficient VitC can induce birth defects through dysregulation of TET1 catalytic activity. However, we also observed divergent phenotypes and extent of DNA hypermethylation at target genes between *Tet1* KO and VitC-deprived embryos, suggesting that TET-independent mechanisms also contribute to the severe embryonic malformations due to VitC deficiency.

The alpha-ketoglutarate dependent dioxygenases are divided up into 4 groups: (i) collagen hydroxylases that stabilize the secondary structure of collagen (*41*), (ii) prolyl hydroxylase containing domain enzymes (PHDs) that mediate the hydroxylation of HIF and are important for oxygen sensing (*42, 43*), (iii) KDM demethylating histones (*44*), and (iv) the TET enzymes (*8, 45*). We ruled out disruption of collagen hydroxylases in VitC-deprived *Gulo* KO embryos. In agreement, others have shown that collagen can be synthesized *in vivo* by *Gulo*^-/-^ mice without a reduction in proline hydroxylation levels, suggesting the existence of VitC-independent collagen synthesis pathways in mice (*29*). Loss of PHD2 (but not PHD1 or PHD3) results in embryonic lethality due to placental and cardiac defects, but the phenotypes are less severe than the ones we observed in *Gulo*^-/-^ (*42*). Moreover, VitC deficient mice do not exhibit impaired HIF-mediated oxygen sensing by PHDs (*46*). Among histone demethylases, KDM3A and KDM3B targeting mono- and di-methylation of histone H3 lysine 9 (H3K9) are the most likely candidates, because they have been shown to be selectively activated by VitC *in vitro* in ESCs and iPSCs, compared to other KDMs (*18, 47*). Further, combined loss of both enzymes leads to severe embryonic malformations around E6.5, indicating that demethylation of H3K9me2 is crucial for embryonic development (*44, 48*). In our Western blot analysis of E11.5 brains from B6-*Gulo* KO mice, we observed that H3K9me2 is the only modification that increased globally in the absence of VitC. In contrast, global levels of H3K4me1, H3K4me3, H3K27me3, and H3K27ac, all important histone PTMs demarcating distinct chromatin states such as active, repressed or bivalent/poised promoters or enhancers, were dramatically lost (*39*). H3K9me2 is one of the most abundant histone PTMs and is associated with heterochromatin organization (*49, 50*). A disruption of steady state H3K9me2 marks by VitC deficiency may have a global impact on higher order chromatin organization that results in the erosion of other epigenetic modifications including DNA methylation. Loss of many histone PTM modifiers (not only KDMs) result in embryonic lethality and gastrulation-associated defects (*12*). We determined the sensitive window of susceptibility to VitC deficiency to occur during gastrulation between E6.5-E8.5, which would be consistent with the disruption of global histone and DNA methylation patterning as the dominant mechanism. In this regard, the DNA hypomethylated state in VitC-deprived E8.5 B6-*Gulo* KO embryos may also involve a secondary effect of histone methylation disruption and dysregulated heterochromatin but otherwise is phenotypically “silent” during gastrulation in the absence of disruptive genetic modifiers. Future investigation should examine whether widespread DNA hypomethylation is coincident with promoter-specific DNA hypermethylation, akin to a cancer and aging epigenome (*51*), or associated with loss of H3K27me3 (and other marks) to mimic a state of epigenome erosion, recently observed in Alzheimer’s disease (*52*). These hypoDMRs are not associated with birth defects but may provide insights into the diseases with higher risks later in life.

Our mouse model of manipulating genetic backgrounds in response to nutritional deficiency shows that inter-strain differences exceeded intra-strain treatment effects in both methylomes and transcriptomes. Strain-dependent methylation variation remains understudied despite its biological relevance. Grimm et al. used WGBS to identify 6,380 strain-specific DMRs between B6 and

C3H/HeN adult mice, mainly at enhancers (*53*). Similarly, Orozco et al. identified 2,865 strain- and allele-specific CpGs using RRBS between B6 and DBA/2J adult mice (*54*). Earlier array-based studies, though limited in coverage, also revealed consistent strain-associated differences (*55, 56*). Targeted reports further highlight locus- and element-specific methylation variability, underscoring the impact of genetic background on the methylome (*57, 58*). We recently mapped a quantitative trait locus (QTL) on chromosome 9 of the 129S6 genome associated with higher risks for NTDs in *Tet1* KO embryos. At the peak center of the QTL is a missense single nucleotide variant (SNV) in *Snx1*, a gene previously implicated in NTDs (*28, 59*). *Snx1* encodes a protein central to endosomal trafficking – the sorting and transport of proteins within cells modulating processes like signaling and endocytosis (*60, 61*). In plants, SNX1 regulates iron homeostasis by recycling the iron transporter IRT1 (*62*); in mammals, its homolog SNX3 mediates transferrin receptor recycling (*63*). Vitamin C functions as a cofactor for dioxygenases by reducing Fe³L to Fe²L at the active site and can stimulate enzyme activity, even in low Fe³L conditions. If Fe²L is abundant, vitamin C becomes less critical (*14, 15*). It is plausible that a deleterious SNV in *Snx1* influences intracellular iron transport, rendering certain strains more reliant on vitamin C’s reducing activity.

VitC is well known to have many physiological functions important for human health and is a cheap readily available supplement. In healthy humans, plasma levels of vitamin C are about 50 μM (*13*). However, in the 2003-2004 National Health and Nutritional Examination survey, over 7% of the US population (>20 million individuals) was found to be VitC deficient, conventionally considered to be having serum ascorbate concentrations <11.4 μM; below this level, there is increased risk of developing scurvy, a condition commonly associated with bleeding gums (*64*). Although overt scurvy has rarely been reported in the US in the past 30 years, milder conditions characterized by fatigue and irritability may be under reported. Marginal VitC deficiency can have serious consequences for fetal brain development, as shown in guinea pig models where prenatal VitC deficiency led to persistent impairment of postnatal hippocampal development which was not alleviated by postnatal repletion (*65*). Additionally, certain pregnancy-related complications, such as hyperemesis gravidarum (HG) (*66*), type I diabetes (*67*) and hypertension (*68*), can lead to nutrient deficiencies, including VitC (*69*). HG, an extreme form of morning sickness leading to dehydration and starvation and common in 1% of pregnancies, presents itself throughout the first trimester of gestation and can lead to abnormal fetal brain development if left untreated (*70*). A study from 1976 found a significant reduction of maternal folate and VitC status in 6 mothers who gave birth to infants with NTDs (*71*). Additionally, higher intakes of vitamin C and E appear to be associated with decreased likelihood of having a pregnancy affected by spina bifida (*72*). However, a landmark vitamin study in 1991 determined folic acid (vitamin B9) supplementation to have a significant (72%) preventive effect against NTDs in previously affected pregnancies, whereas 7 other vitamins including VitC did not (*73*). Since that study, folic acid supplementation and dietary fortification have become the most effective means for preventing birth defects, and hardly any additional studies investigated the impact of VitC deficiencies during pregnancies. Cochrane reviews did not observe any statistically additional health benefits of additional VitC supplementation, which is the stance of the WHO since (*74–76*). Of note, there are still approximately 3,000 NTD-affected pregnancies in the US each year despite folic acid fortification in staple foods (*77*). Our recent study revealed interactions between *Tet1* gene dosage and maternal FA status that suggest that DNA methylation dysregulation by *Tet1* may underlie the resistance of NTDs to FA supplementation (*27*). This raises the question whether promoting timely VitC supplementation by at-risk pregnant mothers could convert folic acid-resistant NTDs to folic acid-responsive NTDs, thus preventing all preventable NTDs.

In the broadest scope, the ramifications of these results extend to all women of reproductive age worldwide. While our primary use of VitC manipulates dioxygenase functions *in utero*, our investigations present a broader model for environmental stressors, providing new insights into the relationship between dynamic changes in the epigenome driven by demethylases, genetic determinants of susceptibility, and environmental exposures that can induce defects in early development. Our results counter the current lack of evidence that VitC supplementation has benefits during pregnancy, by demonstrating the contribution of genetic risk factors, critical exposure periods and optimal dosing. Leveraging VitC’s status as a well-tolerated supplement already approved for daily consumption, our findings hold promise for rapid translation to biomedical applications. Specifically, in the realm of patient-stratified precision medicine, public health management should advocate extra care for patients with nutrient deficiencies during gestation and increase awareness of a correct balance of folic acid with other essential vitamins (*78*).

## Materials and methods

### Mouse breeding and VitC withdrawal

B6-*Gulo*^tm1Mae^ mouse strain, a gift from Sean Morrison (UT Southwestern, TX) (*20*), was either out- crossed for >3 generations to outbred CD1 stocks, or back-crossed for 6-7 generations to the inbred 129S6/SvEvTac (129S6) mice to create an incipient congenic 129S6.B6 strain. Incipient congenic 129S6.B6- *Gulo*^+/-^ mice were intercrossed upon 6-7 backcrossed generations to obtain *Gulo*^-/-^ mice and bred thereafter to homozygosity by filial mating for no more than 3 generations. To maintain the outbred genetic heterogeneity of CD1-*Gulo*^tm1Mae^ mice, we intercrossed heterozygous CD1-*Gulo*^+/-^ only once (F1) during breeding to obtain homozygous CD1-*Gulo*^-/-^, which were mated in experiments with non-sibling offspring from other breeding pairs. Homozygous CD1-*Gulo*^-/-^ were paired with non-sibling heterozygous CD1-*Gulo*^+/-^;*Tet1*^+/-^ to obtain CD1-*Gulo*^-/-^;*Tet1*^+/-^ mice. Due to sub-fertility and runted growth of homozygous *Tet1*^-/-^ mice (*26, 79, 80*), we avoid the interbreeding of *Tet1*^+/-^ mice.

To prevent lethality, *Gulo*^-/-^ mice on all backgrounds are maintained on a custom diet containing stabilized 1% VitC (TD.160301, Envigo Teklad Diets, WI). Pregnant females are further supplemented with 3.3 g/l VitC, given as 3.7 g/L sodium ascorbate (Sigma Cat. No. 11140), in drinking water until pups are weaned. DiTroia *et al*. reported that VitC is depleted within 7-8 days upon VitC withdrawal (*23*). For VitC deprivation experiments (-VitC), *Gulo*^-/-^ dams (7-12 weeks old) were transferred to regular chow (Sniff breeding diet V1124, Germany) without VitC, three days prior to natural timed matings with *Gulo*^-/-^ males (Figure 1a). The morning of finding a copulation plug was considered E0.5. This schedule ensures that at the time of fertilization there is still 50% of serum VitC present to sustain embryo conception, while by E3.5 all VitC is depleted. VitC *Gulo*^-/-^ males were adapted to -VitC conditions only on the morning of the pairing. Pairs were separated on day 4 (7 days after VitC deprivation of the dam). Some experiments were started after 4 days of VitC depletion to test whether the length (in days) of VitC deprivation can be correlated with phenotype severity. All pregnant mice are euthanized by 20 days of VitC deprivation. In the control groups (+VitC), dams receive 3.3 g/l VitC in drinking water during the entire experiment. All experimental procedures on mice were reviewed and approved under project P152/2021 by the KU Leuven Ethical Committee for Animal Experimentation in compliance with the European Directive 2010/63/EU.

### Embryo collection and scoring

Pregnant dams were sacrificed at indicated timepoints using cervical dislocation and dissected to collect both uterine horns. Individual deciduae were separated in a petri-dish containing cold PBS, followed by removal of the outer layers. The embryos were excised from the yolk sac and imaged using a stereo microscope; tissues were dissected if needed and snap frozen in liquid nitrogen or fixed in 4% PFA overnight at 4°C. For embryos ≥ E9.5, part of the yolk sac, visibly free of maternal blood and placental tissues, was collected and used for genotyping. For E8.5 embryos, headfolds were dissected and the remainder of the body was used for genotyping. Genotyping primers are listed in Supplementary Table S1. The phenotype and stage of embryos was preliminarily scored and assigned by a researcher when collecting embryos. These scores were subsequently validated by three independent researchers based on the images. Malformations were classified by whether they were specific to the brain, heart, hind body (scored as “mild”) or affecting multiple organs or the whole embryo grossly (“severe”) (Supplementary Figure 1c). Having two or more types of malformations were also scored as “severe”. When there was no discernible embryonic tissue in the decidua or embryos were disintegrating in the process of being resorbed, we scored them as “resorbing”. Developmental stage of embryos with gestational age ≥ E9.5 was determined using Theiler staging, which uses anatomical characteristics of the embryo(*81*) (Supplementary Figure 1d). When the severity of the malformations precluded stage assignment, embryos were scored as “no Theiler stage” (no TS). E8.5 embryos were staged by somite counts. If there was any uncertainty between the day of fertilization and the timepoint when the embryos were dissected (i.e. due to a missed copulation-plug), the entire litter was excluded from our analysis.

### Plasma measurement of VitC using mass spectrometry

VitC is labile and prone to spontaneous oxidation yielding a mixture of oxidation products: dehydroascorbic acid, 2,3-diketogulonic acid, threonic acid and oxalic acid (*82*). Threonic acid and oxalic acid are the final oxidation products. To obtain a stable and final oxidation product that serves as a proxy for plasma vitamin C concentration, plasma samples were extracted by 70% acetonitrile (Fisher Scientific) at a volume ratio of 1:20 followed by 24-hour incubation at 25°C. After incubation, 2ul of the extract was injected for tandem LC MS/MS measurement of threonic acid. The platform comprised of an Agilent Model 1290 Infinity II liquid chromatography system coupled to an Agilent 6460A Triple Quadrupole MS analyzer. A Dursan® coated Diamond Hydride column (Microsolv) was used for separation. The mobile phases are: (A) 50% isopropanol, containing 0.025% acetic acid (EMD Chemicals), and (B) 90% acetonitrile containing 5 mM ammonium acetate (Sigma-Aldrich). To eliminate the interference of metal ions on chromatographic peak integrity and electrospray ionization, EDTA was added to the mobile phase at a final concentration of 5 µM. The following gradient was applied: 0-1.0 minute (min), 99% B; 1.0-6.0 min, to 2% B; 6.0 to 12.0, 2% B; 12.1 to 17 min, 99% B. LC flow rate is 0.4ml/min and column temperature is 25°C. MRM transitions and acquisition parameters for threonic acid are shown in Supplementary Table S2. Acquired LC/MS/MS data was analyzed by MassHunter Quantitative analysis 10.0 (Agilent Technologies, CA). For absolute quantification, threonic acid standard (Signa-Aldrich) was spiked into plasma samples having undetectable threonic abundance to generate calibration curve.

### Low input whole genome bisulfite sequencing (WGBS) library preparation

400 ng of genomic DNA (gDNA) was sonicated on a Covaris LE220 with settings targeting an average size of 400 bp. Prior to bisulfite conversion, 0.5% unmethylated Lambda gDNA was spiked in and DNA quantification performed using Picogreen. 400 ng of sheared gDNA was bisulfite converted with the Zymo EZ DNA Methylation Gold kit (VWR, D5005) following the product manual. Estimating a 75% recovery rate, 50 ng of deaminated DNA was used for library prep with the IDT xGen™ Methyl-Seq DNA library prep kit (IDT, 10009824). Briefly, gDNA was heat-denatured and the single stranded DNA was used as template for proprietary adaptase stub tailing and adapter ligation prior to full-length adapter addition during the indexing PCR amplification of the library. Following product purification, the resulting libraries were quantitated by Picogreen and fragment size assessed with the Agilent 2100 Bioanalyzer. All samples were pooled equimolarly and re-quantitated by qPCR using the Applied Biosystems ViiA7 Quantitative PCR instrument and a KAPA Library Quant Kit (Roche, KK4824). Using the concentration from the ViiA7™ qPCR machine above, 150 pM of equimolarly pooled library was loaded onto one lane of the NovaSeq S4 flowcell (Illumina, 20028312) following the XP Workflow protocol (Illumina, 20043131) and amplified by exclusion amplification onto a nanowell-designed, patterned flowcell using the Illumina NovaSeq 6000 sequencing instrument. PhiX Control v3 adapter-ligated library (Illumina,/n FC-110-3001) was spiked-in at 2% by weight to ensure balanced diversity and to monitor clustering and sequencing performance. A paired-end 150 bp cycle run was used to sequence an average of 400 million read pairs (∼40X coverage) per sample.

### WGBS analysis

TrimGalore (v0.6.10) was used to trim reads based on quality (PHRED < 30) and to remove adapter sequences. Further, from both read 1 and read 2, 15 bp were trimmed from the 5’ end and 5 bp from the 3’ end. Only reads with a minimum length of 20 bp were kept. Using Bismark (v0.23.1), the trimmed reads were aligned to GENCODE mm10 GRCm38.p6 with a maximal insert size of 500 bp, followed by deduplication and methylation extraction. Using a custom script the CpG counts were merged to one strand. R (v4.0.5) together with the bsseq package (v1.36.0) was used for further downstream analysis. First, CpGs were removed where the coverage was less than the 99.9 percentile per sample, and CpGs were kept that were covered minimally once in each sample. For PCA plots and cumulative distribution plots, samples were merged together, and methylation was calculated over 1kb bins. For DMR analysis, the DSS package (v2.48.0) was used using a standard workflow and parameters (*83*). First smoothing was applied using BSsmooth function implemented in the bsseq framework (*84*). Then, differential CpGs were called where FDR < 0.01 and ΔCpG 5mC > 10%. From these differential CpGs, DMRs were called with FDR < 0.01. DMRs were selected with the following criteria: Δ 5mC > 10%, n CpGs ≥ 3, and length > 100 bp. To prevent false positive DMRs based on coverage differences between conditions, we kept DMRs where the coverage of the DMR was less than 3x the genome wide average per library, and where the total coverage difference between two groups is less than 5x (*53*). All filtering steps were done on unsmoothed data. DMRs were annotated using AnnotatR (v1.16.0) and ChIPseeker (v1.26.2) packages in R (v4.0.5) using UCSC gene feature and CpG island (CpGi) locations. Genes were associated with DMRs using the rGREAT (v1.22.0) package. GO term enrichment was performed using Cluster Profiler (v3.18.1).

### RNA-seq library preparation

RNA was extracted from snap-frozen E8.5 embryonic headfolds using the RNeasy micro kit (Qiagen, 74004), while RNA was extracted from snap-frozen dissected E11.5 embryonic brains using the RNeasy plus mini kit (Qiagen, 74136). RNA purity was assessed by using NanoDrop spectrophotometry (ThermoFisher Scientific), quantified using Picogreen, and RNA integrity was evaluated with a 2100 Bioanalyzer and RNA chips (Agilent Technologies, Santa Clara, CA). E8.5 headfold cDNA libraries were prepared from 10 ng of total RNA using the SMARTer® Stranded Total RNA-Seq v3 - Pico Input Mammalian kit (Takara, 634485). E11.5 brain RNA-seq libraries were prepared with 250 ng of input material, using the TruSeq Stranded mRNA Library Prep kit (Illumina, 20020594), according to the according to the manufacturer’s protocol. Library quantification and pooling were performed equimolarly using the KAPA Library Quantification kit for Illumina platforms (KAPA Biosystems, Wilmington, MA). Sequencing was conducted on the Illumina NovaSeq 6000 platform, generating approximately 60 million read pairs per sample for low input samples, and 20 million read pairs for regular polyA-selected libraries.

### RNA-seq analysis

RNA-seq analysis followed a general processing pipeline: Using Cutadapt (v4.9) (*85*), polyA/T tails and bad quality reads were trimmed, followed by two more rounds of trimming to remove R1 and R2 adapters `(-a "r1adapter=AGATCGGAAGAGC" -A "r2adapter=AGATCGGAAGAGC"`) first with an overlap of minimally 3 and an max error rate of 0.1, and next a minimal overlap of 20. Reads were aligned to the genome, GENCODE mm10 GRCm38.p6, using STAR aligner (v2.7.11a) (*86*). Since the low input RNA-seq libraries contain a unique molecular identifier, UMI-tools was used for deduplication of these (*87*). For both library types, featureCounts (v2.0.6) (*88*) was used to count reads and rsem (v1.3.1) (*89*) was used to calculate TPM values. Subsequently, the counts were imported into R (v4.0.2) and differentially expressed genes were defined using DEseq2 (v1.30.0) (*90*) (FDR adjusted p.val < 0.05) and log fold changes corrected using “ashr” method (*91*). GO term enrichment was performed using Cluster Profiler (3.18.1) (*92*).

### Targeted bisulfite-sequencing

Targeted amplicon bisulfite sequencing was performed as described previously (*11, 27*). We assayed loci at 3 gene promoters by designing primers to detect on average 2 amplicons of 250-280 bp each at each locus. Genomic DNA (gDNA) was extracted from E11.5 brains using the Purelink genomic DNA Mini Kit (Invitrogen, K182001), according to the manufacturer’s instructions. The quality of the gDNA was assessed using Nanodrop and the absence of RNA contamination was verified by running the samples on a 0.8% agarose gel stained with SyberSafe. 1.5 µg of gDNA was used for bisulfite conversion, using the EpiTect Fast DNA Bisulfite kit (Qiagen, 59824) and eluted in 15 µl of elution buffer provided in the kit. 0.5 µl of bisulfite converted gDNA was used in a 20 ul PCR reaction containing 300 nM each of a forward and a reverse primer containing P7 and P5 tails respectively, Platinum^TM^ *Taq* DNA polymerase High Fidelity (Invitrogen, 11304-011) and kit buffers. Oligos are listed in Supplementary Table S1. PCR products were loaded on a 1.5% agarose gel for gel extraction of individual amplicon bands of interest using PureLink Quick Gel Extraction kit (Invitrogen, K210012). The concentration of each amplicon was measured using Qubit^TM^ dsDNA HS Assay kit (Invitrogen, Q32854) and diluted to 15 nM. The quality of the pooled amplicons was assessed using fragment analyzer (Agilent) and the Qubit™ dsDNA HS Assay kit (Invitrogen, Q32854). The amplicon pools were diluted to max 5 ng/µl and combined for a secondary PCR to add indexes and sequencing adapters. The PCR reaction was set up as follows: 9 µl DNA, 0.5 µl custom p7 primer (125 nM), 0.5 µl custom p5 primer (125 nM) and 1x 10 µl Phusion® High Fidelity PCR master Mix with HF buffer (Biolabs new England M0531S). The following program was used in a thermocycler: 94 °C 30 seconds (sec); 15x 94 °C 10 sec, 51 °C 30 sec, 72 °C 30 sec; 72 °C 1 min. The custom primers are provided with unique dual indexes to label the samples. The resulting library was purified with a 1x clean-up using AMPure XP beads following manufacturers protocol. The final quality of libraries was analyzed using fragment analyzer (Agilent) and pooled equimolar. The concentration of the final pool was measured using qPCR (Kapa SYBR fast, Roche, KK4600) and loaded on a NovaSeq for PE150 sequencing for a minimum of 200,000 reads per amplicon and on average 350,000. Using Trim Galore! (v0.6.7) , reads were trimmed based on quality (PHRED < 20), adapters were removed and reads of fewer than 20 bp were excluded. Using Bismark (v0.23.1), the trimmed reads were aligned to GENCODE mm10 GRCm38.p6) with a maximal insert size of 500 bp, followed by methylation extraction. Only CpGs with a minimal coverage of 1000x were retained for analysis and plotted over the genomic locus assayed using a custom script in R (v4.0.3).

### Picrosirus red staining

E11.5 CD1 embryos were fixed in 4% paraformaldehyde (PFA) in PBS (Gilbo, 10010023) and stored in 70% ethanol in saline (0.9% NaCl). Severely malformed embryos were pre-embedded in 1.5% UltraPureTM Agarose (Invitrogen, 16500500) in saline to achieve the desired embryo orientation. For this, embryos were rinsed twice in saline and twice in 1.5% UltraPureTM Agarose (Invitrogen, 16500500) at 60°C. Embryos were positioned under microscopy and gel blocks containing the embryos were cut using a scalpel once the agarose was set. The agarose blocks and the remaining embryos were then transferred to a cassette and stored at 4°C in 70% ethanol. Thereafter, tissue processing (dehydration) was performed by the histology core facility of the Centre for Molecular and Vascular Biology (CMVB, KU Leuven). After dehydration, embryos were embedded in paraffin, sectioned (5 µm thickness) with a microtome, and dried on a hot plate at 37°C. For staining, sections were deparaffinized by 2 × 10 min immersions in xylene. Rehydration was performed by dipping the sections in decreasing concentrations of ethanol (100%, 95%, 90%, 80%, 70%, 60%, and 50%) for 2 min at each concentration, followed by immersion for 5 min in water at RT. Sections were stained for collagen using the Picrosirius red solution (Vitro ViewTM Picro-Sirius Red Stain Kit,GeneCopoeia, VB-3017) for 1 hour, washed in two changes of acidified water (0.5% glacial acetic acid), washed twice in deionized water, dehydrated in 90%, 95%, and 100% ethanol for 2 min each step, and finally cleared in 2 changes of xylene for 5 min each. To counter-stain nuclei, an 8 min immersion in Weigert’s haematoxylin was performed, followed by washing for 10 min in running tap water. Stained sections were mounted in a resinous medium and imaged using a Axiovert 40 CFL inverted microscope (Carl Zeiss).

### Western blotting

Snap frozen mouse E11.5 brains were lysed using ice-cold high-salt RIPA buffer (50 mM Tris at pH 8.0, 600 mM NaCl, 0.2 mM EDTA, 1% NP-40, 0.5% sodium deoxycholate, 0.1% SDS) containing 1 mM phenylmethylsulfonyl fluoride, 0.5 mM DTT, phosphatase inhibitor cocktail 2 and 3 (Sigma-Aldrich, P5726 and P0044) and protease inhibitor cocktail (Roche, 11836153001), and further disrupted ice-cold using a T10 basic tissue homogenizer (IKA) for 10 minutes to lyse the tissues. Samples were sonicated in a Bioruptor (Diagenode) for 5 cycles (30sec on, 30sec off) on high at 4°C to fragment all chromatin, centrifuged for 15 min at 16.000 r.c.f. at 4°C, following which the supernatant was collected and stored at -80°C. Protein concentration was measured using Bradford assay in a 96-well micro plate format. Protein samples were prepared in 1x Laemmli sample buffer (62.5 mM Tris-HCl at pH 6.8, 2.5% SDS, 0.002% bromophenol blue, 5% β-mercaptoethanol, 10% glycerol) by boiling for 10 min at 95°C. 20 µg of protein was loaded on an 12% SDS–polyacrylamide gel. Samples were run in 1X running buffer (25mM Tris, 192 mM glycine, 0.1% SDS) and then transferred to a PVDF membrane with transfer buffer (25 mM Tris, 192 mM glycine and 20% methanol). Membranes were blocked with 5% non-fat dry-milk Tris-buffered saline with 0.1% Tween- 20 (TBS-T), incubated overnight at 4°C with primary antibodies diluted in 5% non-fat milk, and subsequently with HRP-conjugated secondary antibodies diluted 1:5000 in TBS-T with 5% non-fat milk for 1 h at room temperature. The signal was detected using Clarity Western ECL substrate (Bio-Rad 1705060) on AGFA Curix 60 Film Processor. Primary antibodies used in this study are: anti-ACTB (Cell-Signaling, 4970, 1:1000), anti-H3 (Abcam, ab1791, 1:5000), anti-H3K27me3 (Upstate, 07-449, 1:1000), anti-H3K4me3 (Abcam, ab8580,1:1000), anti-H3K4me1(EpiCypher, 13-0057, 1:1000), anti-H3K9me2 (Actif-Motif, 39239, 1:5000), anti-H3K9me3 (Active Motif, 39766, 1:1000) and anti-H3K27ac (Actif-Motif, 39135, 1:1000).

### Data Availability

All WGBS and RNA-seq data is available at https://www.ncbi.nlm.nih.gov/geo/ With accession number GSE295926.

To review GEO accession GSE295926:

Go to https://www.ncbi.nlm.nih.gov/geo/query/acc.cgi?acc=GSE295926

Enter token gjivywcmjzsdbiv into the box

## Acknowledgement

C57Bl/6-*Gulo*^tm1Mae^ were kindly provided by Dr. Sean Morrison, Children’s Medical Center Research Institute at UT Southwestern, Dallas, Texas, USA. We thank Luís Pereira Pina and Mariana Schroiff for technical assistance with mouse experimentation and tissue collection. Low input WGBS, Takara SMARTer stranded low-input total RNA-seq, and poly A-selected RNA-seq library preparations and sequencing were performed by Ping Kang and Daniel Kraushaar at the BCM GARP Core.

## Author contributions

K.P.K and B.K.V. designed and conceived the study. K.P.K. directed and supervised the study. K.P.K. and B.K.V. interpreted the data. B.K.V. performed or supervised all experiments and molecular analysis techniques. C.C. and W.B. maintained mice colony breeding, performed VitC dosing studies and collected embryos and plasma. R.C. performed embryo dissections, targeted bisulfite sequencing and western blotting. S.C.T. performed Sirus red staining. H.C., Q.C. and S.S.G. performed LC-MS/MS measurement of plasma ascorbate. B.T and K.D.R. provided expertise on methylome analysis. D.K provided expertise on low-input WGBS and RNA-seq library preparations. R.H.F. provided expertise on the analysis of quantitative trait loci and NTDs. K.P.K., B.K.V., and R.C. validated all embryo phenotyping scores. B.K.V. performed WGBS and RNA-seq analysis. K.P.K and B.K.V. wrote the manuscript and prepared the figures.

## Supplementary Materials

Supplementary Figure 1 (related to main Figure 1)

**a**, Plasma levels of total ascorbate detected as the degradation product threnoic acid by LC-MS/MS in B6-*Gulo*^-/-^ and 129S6.B6*Gulo*^-/-^ female mice at the indicated timepoints of VitC withdrawal, compared to WT mice per strain. d0, dam maintained on 1% stabilized VitC diet prior to VitC withdrawal. The gestational time corresponding to days of maternal VitC withdrawal assumes copulation within the first day of pairing with a stud male, i.e. day 4 after VitC withdrawal. **b**, Three representative +VitC *Gulo*^-/-^ embryos and a representative full litter of -VitC *Gulo*^-/-^ embryos of each background. **c**,**d**, Representative images of malformations (c) and delays (d) observed in VitC-deprived embryos. Overview images in (b-d) are assembled from individual embryo pictures. Scale bars indicate 1 mm. **e,** Representative images of hemorrhagic placentas in -VitC CD-*Gulo*^-/-^ and 129S6.B6-*Gulo*^-/-^. **f,** Phenotype penetrance scores of -VitC B6, CD1, and 129S6.B6 *Gulo*^-/-^ embryos, stratified by days of VitC depletion before detection of copulation plug, i.e. fertilization. Chi-square test was used for statistics. “Mild malformation” (malf.) = 1 malformation, “severe malformation” = multiple malformations or grossly malformed. “Resorbing” = empty decidua or embryos in the process of being resorbed. “no TS” = not possible to assign a Theiler stage. n = number of embryos per condition, litters (lit.) = number of litters per condition. Statistical comparisons are made using Chi-square test. **g**, Number of resorptions per litter, scored as incidences of resorbing embryos in the process being resorbed or empty decidua. **h**, Number of embryos per genetic background and treatment. Pairwise comparisons are done using Mann-Whitney U test.

Supplementary Figure 2 (related to main Figure 1)

**a**, Sirius red staining (red = collagen) in +VitC (i, ii) and -VitC (iii, iv) CD1-*Gulo*^-/-^ embryos. **i**, **ii**, **iv**, are embryos scored as “normal”, while **iii** is an embryo with severe malformations “affected” by VitC deprivation. Scale bars in the insets indicate 100 µM, while scale bars in the whole embryo images indicate 0.5 mm. **b**, **c**, Representative images of E8.5 (b, left) and E9.5 (c, left) +VitC and -VitC *Gulo*^-/-^ embryos and associated staging scores (right) in B6, CD1, and 129S6.B6 genetic backgrounds at E8.5 (b) and in CD1 background at E9.5 (**c**). The scale bar is 0.5 mm for E8.5 and 1 mm for E9.5 images. E8.5 embryos were staged by somite counts, not Theiler stage. “mild malformation” (malf.) = 1 malformation, “severe malformation” = multiple malformations or grossly malformed, “Resorbing” = empty decidua or embryos in the process of being resorbed, “no TS” = not possible to assign a Theiler stage, n = number of embryos per condition, litters (lit.) = number of litters per condition. Pairwise comparisons were performed using Chi-square test.

Supplementary Figure 3 (related to main Figure 2)

**a**, Images of the individual E8.5 embryos collected for WGBS. **b**, Table listing the experimental identifiers, sex, somite counts of individual embryos used in WGBS and the sample library coverage and global non-CpG methylation levels. **c**, **d**, Duplication rate (**c**) and global CpG methylation levels

(**d**) of WGBS libraries. **e**, PCA of CpG methylomes in all samples grouped by strains (B6 and 129S6.B6), treatment (-VitC and +VitC) and sex (male and female). The two 129S6.B6 -VitC pheno samples are separately shown in black color. **f, g,** Global CpG methylation levels at indicated genomic elements per library from *Gulo*^-/-^ headfolds of B6 (f) and 129S6.B6 (g) strains. **h**, Enrichment of 129S6.B6 DMRs overlapping with functional genomic elements active during mouse fetal development (timepoints are indicated) as defined in ENCODE3. The 15 chromatin states shown in Figure 2h condensed into 5 classes: En, enhancers; Pr, promoters; Tr, transcription; Hc, heterochromatin and NS, no signal. **i**, Volcano plot of differentially methylated CpGs, highlighted in orange and grey indicating those with significant gain or loss, respectively, in -VitC relative to +VitC, (left), histogram of DMR distribution by differences in CpG methylation levels in intervals of 10% (right), and number of hyper and hypo DMRs (bottom) in the two outlier 129S6.B6-*Gulo*^-/-^ -VitC pheno headfolds compared to +VitC controls (n=4). **j**, Distribution of hyper- and hypo- DMRs by proximity to CpG islands (CpGi) in the two 129S6.B6-*Gulo*^-/-^ -VitC pheno headfolds.

Supplementary Figure 4 (related to main Figure 2)

**a**, Table listing the experimental identifiers, sex and somite counts of E8.5 embryos used in RNA-sequencing. **b**, PCA plot of all RNA-seq data from all samples showing an outlier TP155.5 (B6,

+VitC, female). **c**, PCA plot of transcriptomes after exclusion of outlier TP155.5. **d**, Number of RNA-seq DEGs defined by pairwise comparison of -VitC versus +VitC treatments per *Gulo*^-/-^ strain, classified by up or downregulation relative to +VitC. **e**, Cumulative distribution of the distance to the nearest B6 hypo-DMR among up- and down-regulated DEGs in B6-*Gulo*^-/-^ headfolds. **f,** Top 10 GO terms associated with up- (left) and down-regulated (right) DEGs in E8.5 B6-*Gulo*^-/-^ headfolds. **g**, Top 10 GO terms associated with down-regulated DEGs in 129S6.B6-*Gulo*^-/-^ headfolds.

Supplementary Figure 5 (related to main Figure 3)

**a**, Table listing the experimental identifiers, phenotypes, developmental stages of the individual embryos used for WGBS and the sample library coverage and global non-CpG methylation levels. **b**, PCA plot of WGBS data from E11.5 brains, collected from each VitC dose exposure from 3300-0 mg/l. **c**, Volcano plots of differentially methylated CpGs, highlighted in orange and grey to indicate those with significant gain or loss, respectively, and histogram of DMR distribution by differences in CpG methylation levels in intervals of 10% (right) in pairwise comparisons of morphologically normal E11.5 brains from each VitC dose exposure at 330 ml/l, 100 mg/l and 33 mg/l compared to +VitC (3300 mg/l) control (left column), in completely VitC-deprived and deformed (0 mg/l, -VitC) compared to +VitC control brains and in phenotypically affected brains compared to morphologically normal brains at 100 mg/l and 33 mg/l (right column). **d**, Global CpG methylation levels within genomic features associated with CpGi, gene bodies and retrotransposable repeats across the genome in staged-matched morphologically normal E11.5 brains exposed to dose-titration of maternal VitC (orange to grey colouring) and those with phenotypes (blue). **e**, Distribution of DMRs induced by half logarithmic dose reduction in VitC until 33 mg/l in morphologically normal E11.5 brains and in completely VitC deprived brain, per pairwise comparison relative to +VitC control, at gene body elements. **f**, **g**, Distribution of hyperDMRs and hypoDMRs associated with NTD or growth delay versus morphologically normal brains per VitC dose by CpG island proximity (f) and genetic elements. **h-i**, Enrichment of NTD-associated hyper-DMRs at 100 mg/l VitC treatment at ENCODE3 genomic elements active during mouse fetal development from E11.5 to P0 (h) and the top 10 enriched GO terms overlapping with ENCODE3 enhancers (i). **j**, **k**, Enrichment of growth delay- associated hyper-DMRs at 33 mg/l VitC treatment at ENCODE3 genomic elements (j) and the top 10 enriched GO terms overlapping with ENCODE3 enhancers (k). Refer to genome reference of ENCODE3 functional elements and legends in Supplementary Figure 3h.

Supplementary Figure 6 (related to main Figure 3)

**a**, Table listing the experimental identifiers, sex, phenotype and stage of the individual E11.5 CD1- *Gulo*^-/-^ embryos used for RNA-sequencing. **b**, Heatmap with hierarchical clustering based on sample-to-sample distances of samples with VitC dose treatment from 3300-0 mg/l and rescue at E6.5. **c**, Number of RNA-seq DEGs defined by pairwise comparison of each indicated VitC dose treatment vs +VitC, classified by up or downregulation. **d**, Number of RNA-seq DEGs defined by pairwise comparison of 100 mg/l -NTD vs 100 mg/l (left) and 33 mg/l -del vs 33 mg/l (right), classified by up or downregulation.

Supplementary Figure 7 (related to main Figure 3)

**a**, Venn diagram showing overlap of hyperDMRs from all pairwise comparisons of each VitC dose treatment versus +VitC control to define *de novo* hyperDMRs (circled in color) that accrue with each half logarithmic dose reduction and stayed hypermethylated at subsequently lower doses. **b**, Dose-response of CpG methylation levels in subsets of *de novo* hyperDMRs induced by each titrated VitC dose reduction from 3300 to 0 mg/l. The box represents the interquartile range (IQR) and the line within indicates the median. **c**, Enrichment of *de novo* hyper-DMRs at 33 mg/l and 0 mg/l VitC dose treatments at ENCODE3 genomic elements active throughout mouse fetal development. **d**, Top 10 enriched GO terms of 33 mg/l *de novo* hyperDMRs that are overlapping with ENCODE3 enhancers. **e**, Top 10 enriched GO terms of 0 mg/l *de novo* hyperDMRs that are overlapping with ENCODE3 enhancers.

Supplementary Figure 8 (related to main Figure 4)

**a**, Volcano plot of differentially methylated CpGs, highlighted in orange and grey to indicate those with significant gain or loss, respectively, in an E11.5 CD1-*Gulo*^-/-^ brain fully rescued by high dose VitC re-supplementation at E6.5 relative to +VitC brain (left), histogram of DMR distribution by differences in CpG methylation levels in intervals of 10% (right), and number of hyper and hypo DMRs (bottom) in **b**, **c,** Distribution of hyperDMRs and hypoDMRs by CpG island proximity (b) and gene body elements (c) in E11.5 brains after re-supplementation at E6.5. **d**, Top 10 enriched GO terms of resistant hyper hyperDMRs resistant to VitC re-supplementatio, as defined in Figure 4f. **e,** Number of RNA-seq DEGs defined by pairwise comparison of a VitC-deprived E11.5 brains (n=2) rescued at E6.5 vs VitC+ control brains (n=2). **f**, Top 10 GO terms associated down-regulated DEGs defined by pairwise comparison of VitC-deprived brains rescued at E6.5 vs VitC+ controls.

Supplementary Figure 9 **a**, **b**, CpG methylation levels at the promoters of *Cspg4*, *Gal*, and *Ngb* in E11.5 brains collected from +VitC and -VitC B6 (**a**) and 129S6.B6 (**b**) *Gulo*^-/-^ embryos. n = 2 individual embryos for +VitC, n = 6 individual embryos for -VitC for both B6 and 129S6.B6. Pairwise statistical comparison is made using a paired t-test. **c**, Images of actual embryos used for targeted bisulfite sequencing in **a**, **b**, from +VitC and -VitC 129S6.B6-*Gulo*^-/-^ embryos. We collected the entire brain, consisting of the hindbrain (HB), midbrain (MB), and forebrain (FB). Dotted lines indicate dissection planes. The image is assembled from individual embryo pictures. **d**, Non-weighted average CpG methylation per amplicon, per individual E11.5 brain collected from +VitC and -VitC B6 and 129S6.B6-Gulo-/- embryos. Dark grey denotes samples in which whole embryos were collected because of gross malformations. **e**, Western blot showing global H3K9me3, H3K9me2, H3K27me3, H3K27ac, H3K4me3, H3K4me1 content in E11.5 brains collected from +VitC and -VitC B6-*Gulo*^-/-^ embryos. As a control, a B6-*Gulo*^+/+^ is also included. H3 and ACTB are used as loading controls. **f**, ImageJ densitometric measurement of each histone mark in -VitC samples relative to VitC-replete controls.

---

**Supplementary Table S1.**
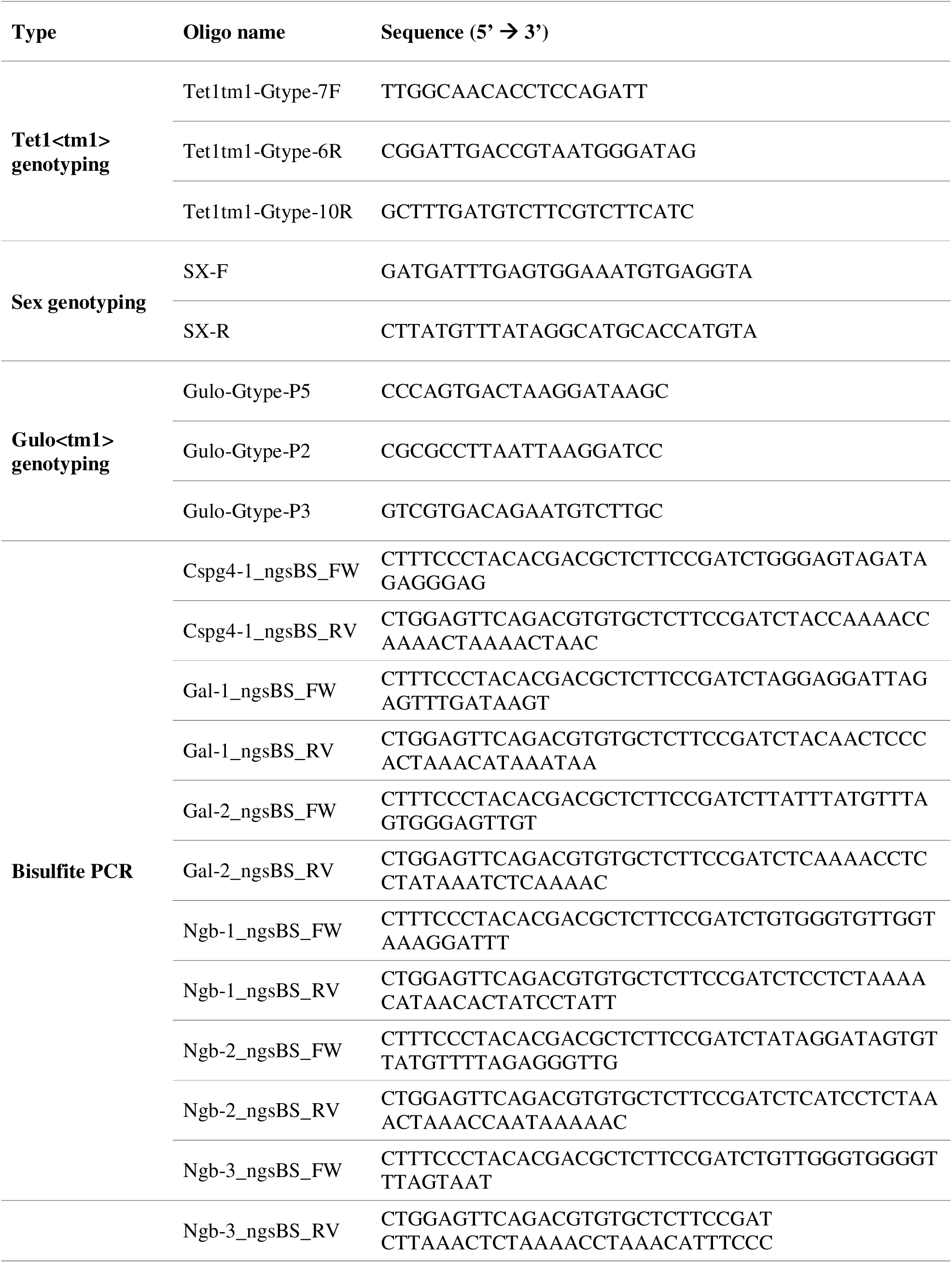
Genotyping and targeted bisulfite-sequencing primers.

**Supplementary Table S2.** LC MS/MS MRM transitions for threonic acid measurement.

| Cpd Name | Prec Ion | Prod Ion | Frag (V) | CE (V) | Cell Acc (V) | Ret Time (min) | Ret Window | Polarity |
| --- | --- | --- | --- | --- | --- | --- | --- | --- |
| Threonic acid | 135.02 | 117 | 68 | 6 | 7 | 5 | 5 | Negative |
| Threonic acid | 135.02 | 75 | 68 | 6 | 7 | 5 | 5 | Negative |
| Threonic acid | 135.02 | 73 | 68 | 6 | 7 | 5 | 5 | Negative |

## References

1. T. P. Fleming et al., Origins of lifetime health around the time of conception: causes and consequences. Lancet 391, 1842–1852 (2018).

2. S. Li, M. Chen, Y. Li, T. O. Tollefsbol, Prenatal epigenetics diets play protective roles against environmental pollution. Clin Epigenetics 11, 82 (2019).

3. , S. National Center for Health, Ed. (Hyattsville, MD, 2022), vol. 71.

4. M. Wadman, United Kingdom moves to prevent birth defects. Science 374, 518–519 (2021).

5. C. Johansson et al., The roles of Jumonji-type oxygenases in human disease. Epigenomics 6, 89–120 (2014).

6. S. M. Kooistra, K. Helin, Molecular mechanisms and potential functions of histone demethylases. Nat Rev Mol Cell Biol 13, 297–311 (2012).

7. S. C. Wu, Y. Zhang, Active DNA demethylation: many roads lead to Rome. Nat Rev Mol Cell Biol 11, 607–620 (2010).

8. S. Ito et al., Tet proteins can convert 5-methylcytosine to 5-formylcytosine and 5- carboxylcytosine. Science 333, 1300–1303 (2011).

9. R. Argelaguet et al., Multi-omics profiling of mouse gastrulation at single-cell resolution. Nature 576, 487–491 (2019).

10. O. Bogdanovic et al., Active DNA demethylation at enhancers during the vertebrate phylotypic period. Nat Genet 48, 417–426 (2016).

11. B. K. van der Veer et al., Dual functions of TET1 in germ layer lineage bifurcation distinguished by genomic context and dependence on 5-methylcytosine oxidation. Nucleic Acids Res 51, 5469–5498 (2023).

12. S. Grosswendt et al., Epigenetic regulator function through mouse gastrulation. Nature 584, 102–108 (2020).

13. J. I. Young, S. Zuchner, G. Wang, Regulation of the epigenome by vitamin C. Annu Rev Nutr 35, 545–564 (2015).

14. C. Loenarz, C. J. Schofield, Physiological and biochemical aspects of hydroxylations and demethylations catalyzed by human 2-oxoglutarate oxygenases. Trends Biochem Sci 36, 7–18 (2011).

15. A. Monfort, A. Wutz, Breathing-in epigenetic change with vitamin C. EMBO Rep 14, 337–346 (2013).

16. K. Blaschke et al., Vitamin C induces Tet-dependent DNA demethylation and a blastocyst-like state in ES cells. Nature 500, 222–226 (2013).

17. J. Chen et al., Vitamin C modulates TET1 function during somatic cell reprogramming. Nat Genet 45, 1504–1509 (2013).

18. J. Chen et al., H3K9 methylation is a barrier during somatic cell reprogramming into iPSCs. Nat Genet 45, 34–42 (2013).

19. J. P. Brabson, T. Leesang, S. Mohammad, L. Cimmino, Epigenetic Regulation of Genomic Stability by Vitamin C. Front Genet 12, 675780 (2021).

20. N. Maeda et al., Aortic wall damage in mice unable to synthesize ascorbic acid. Proc Natl Acad Sci U S A 97, 841–846 (2000).

21. H. Kim et al., Vitamin C deficiency causes severe defects in the development of the neonatal cerebellum and in the motor behaviors of Gulo(-/-) mice. Antioxid Redox Signal 23, 1270–1283 (2015).

22. Y. Kishimoto et al., Insufficient ascorbic acid intake during gestation induces abnormal cardiac dilation in fetal and neonatal SMP30/GNL knockout mice. Pediatr Res 73, 578–584 (2013).

23. S. P. DiTroia et al., Maternal vitamin C regulates reprogramming of DNA methylation and germline development. Nature 573, 271–275 (2019).

24. M. Agathocleous et al., Ascorbate regulates haematopoietic stem cell function and leukaemogenesis. Nature 549, 476–481 (2017).

25. L. Cimmino et al., Restoration of TET2 function blocks aberrant self-renewal and leukemia progression. Cell 170, 1079–1095 e1020 (2017).

26. R. Khoueiry et al., Lineage-specific functions of TET1 in the postimplantation mouse embryo. Nat Genet 49, 1061–1072 (2017).

27. L. Chen et al., The DNA demethylase TET1 modifies the impact of maternal folic acid status on embryonic brain development. EMBO Rep 26, 175–199 (2025).

28. B. K. v. d. Veer, et al., Epigenetic regulation by TET1 in gene-environmental interactions influencing susceptibility to congenital malformations. bioRxiv, 2024.2002.2021.581196 (2024).

29. K. K. Parsons, N. Maeda, M. Yamauchi, A. J. Banes, B. H. Koller, Ascorbic acid-independent synthesis of collagen in mice. Am J Physiol Endocrinol Metab 290, E1131–1139 (2006).

30. D. U. Gorkin et al., An atlas of dynamic chromatin landscapes in mouse fetal development. Nature 583, 744–751 (2020).

31. E. P. Consortium et al., Expanded encyclopaedias of DNA elements in the human and mouse genomes. Nature 583, 699–710 (2020).

32. H. Schorle, P. Meier, M. Buchert, R. Jaenisch, P. J. Mitchell, Transcription factor AP-2 essential for cranial closure and craniofacial development. Nature 381, 235–238 (1996).

33. H. Kim et al., The analysis of vitamin C concentration in organs of gulo(-/-) mice upon vitamin C withdrawal. Immune Netw 12, 18–26 (2012).

34. S. Chrysanthou et al., The DNA dioxygenase Tet1 regulates H3K27 modification and embryonic stem cell biology independent of its catalytic activity. Nucleic Acids Res 50, 3169–3189 (2022).

35. P. Stolz et al., TET1 regulates gene expression and repression of endogenous retroviruses independent of DNA demethylation. Nucleic Acids Res 50, 8491–8511 (2022).

36. R. R. Zhang et al., Tet1 regulates adult hippocampal neurogenesis and cognition. Cell Stem Cell 13, 237–245 (2013).

37. A. Rudenko et al., Tet1 is critical for neuronal activity-regulated gene expression and memory extinction. Neuron 79, 1109–1122 (2013).

38. A. J. Towers, et al., Epigenetic dysregulation of Oxtr in Tet1-deficient mice has implications for neuropsychiatric disorders. JCI Insight 3, (2018).

39. Z. Dai, V. Ramesh, J. W. Locasale, The evolving metabolic landscape of chromatin biology and epigenetics. Nat Rev Genet 21, 737–753 (2020).

40. T. A. Macrae, J. Fothergill-Robinson, M. Ramalho-Santos, Regulation, functions and transmission of bivalent chromatin during mammalian development. Nat Rev Mol Cell Biol, (2022).

41. P. Rappu, A. M. Salo, J. Myllyharju, J. Heino, Role of prolyl hydroxylation in the molecular interactions of collagens. Essays Biochem 63, 325–335 (2019).

42. K. Takeda et al., Placental but not heart defects are associated with elevated hypoxia-inducible factor alpha levels in mice lacking prolyl hydroxylase domain protein 2. Mol Cell Biol 26, 8336–8346 (2006).

43. C. J. Schofield, P. J. Ratcliffe, Oxygen sensing by HIF hydroxylases. Nat Rev Mol Cell Biol 5, 343–354 (2004).

44. E. Dimitrova, A. H. Turberfield, R. J. Klose, Histone demethylases in chromatin biology and beyond. EMBO Rep 16, 1620–1639 (2015).

45. M. Tahiliani et al., Conversion of 5-methylcytosine to 5-hydroxymethylcytosine in mammalian DNA by MLL partner TET1. Science 324, 930–935 (2009).

46. K. J. Nytko et al., Vitamin C is dispensable for oxygen sensing in vivo. Blood 117, 5485–5493 (2011).

47. K. T. Ebata et al., Vitamin C induces specific demethylation of H3K9me2 in mouse embryonic stem cells via Kdm3a/b. Epigenetics Chromatin 10, 36 (2017).

48. S. Kuroki et al., Combined Loss of JMJD1A and JMJD1B Reveals Critical Roles for H3K9 Demethylation in the Maintenance of Embryonic Stem Cells and Early Embryogenesis. Stem Cell Reports 10, 1340–1354 (2018).

49. Q. Guo, S. Sidoli, B. A. Garcia, X. Zhao, Assessment of Quantification Precision of Histone Post-Translational Modifications by Using an Ion Trap and down To 50L000 Cells as Starting Material. J Proteome Res 17, 234–242 (2018).

50. A. Poleshko et al., H3K9me2 orchestrates inheritance of spatial positioning of peripheral heterochromatin through mitosis. Elife 8, (2019).

51. J. Campisi, Aging, cellular senescence, and cancer. Annu Rev Physiol 75, 685–705 (2013).

52. X. Xiong et al., Epigenomic dissection of Alzheimer’s disease pinpoints causal variants and reveals epigenome erosion. Cell 186, 4422–4437 e4421 (2023).

53. S. A. Grimm et al., DNA methylation in mice is influenced by genetics as well as sex and life experience. Nat Commun 10, 305 (2019).

54. L. D. Orozco et al., Intergenerational genomic DNA methylation patterns in mouse hybrid strains. Genome Biol 15, R68 (2014).

55. H. Gujar, J. W. Liang, N. C. Wong, K. Mozhui, Profiling DNA methylation differences between inbred mouse strains on the Illumina Human Infinium MethylationEPIC microarray. PLoS One 13, e0193496 (2018).

56. E. Schilling, C. El Chartouni, M. Rehli, Allele-specific DNA methylation in mouse strains is mainly determined by cis-acting sequences. Genome Res 19, 2028–2035 (2009).

57. K. Padjen, S. Ratnam, U. Storb, DNA methylation precedes chromatin modifications under the influence of the strain-specific modifier Ssm1. Mol Cell Biol 25, 4782–4791 (2005).

58. S. Wang et al., Strain differences between CD-1 and C57BL/6 mice in expression of metabolic enzymes and DNA methylation modifications of the primary hepatocytes. Toxicology 412, 19–28 (2019).

59. D. G. Schwarz, C. T. Griffin, E. A. Schneider, D. Yee, T. Magnuson, Genetic analysis of sorting nexins 1 and 2 reveals a redundant and essential function in mice. Mol Biol Cell 13, 3588–3600 (2002).

60. N. Vieira, T. Rito, M. Correia-Neves, N. Sousa, Sorting Out Sorting Nexins Functions in the Nervous System in Health and Disease. Mol Neurobiol 58, 4070–4106 (2021).

61. B. Zheng et al., Essential role of RGS-PX1/sorting nexin 13 in mouse development and regulation of endocytosis dynamics. Proc Natl Acad Sci U S A 103, 16776–16781 (2006).

62. T. Brumbarova, R. Ivanov, SNX1-mediated protein recycling: Piecing together the tissue-specific regulation of arabidopsis iron acquisition. Plant Signal Behav 13, e1411451 (2018).

63. C. Chen et al., Snx3 regulates recycling of the transferrin receptor and iron assimilation. Cell Metab 17, 343–352 (2013).

64. R. L. Schleicher, M. D. Carroll, E. S. Ford, D. A. Lacher, Serum vitamin C and the prevalence of vitamin C deficiency in the United States: 2003-2004 National Health and Nutrition Examination Survey (NHANES). Am J Clin Nutr 90, 1252–1263 (2009).

65. P. Tveden-Nyborg et al., Maternal vitamin C deficiency during pregnancy persistently impairs hippocampal neurogenesis in offspring of guinea pigs. PLoS One 7, e48488 (2012).

66. J. M. Roberts et al., Vitamins C and E to prevent complications of pregnancy-associated hypertension. N Engl J Med 362, 1282–1291 (2010).

67. B. Juhl, F. F. Lauszus, J. Lykkesfeldt, Poor Vitamin C Status Late in Pregnancy Is Associated with Increased Risk of Complications in Type 1 Diabetic Women: A Cross-Sectional Study. Nutrients 9, (2017).

68. V. London, S. Grube, D. M. Sherer, O. Abulafia, Hyperemesis Gravidarum: A Review of Recent Literature. Pharmacology 100, 161–171 (2017).

69. M. E. van Stuijvenberg, I. Schabort, D. Labadarios, J. T. Nel, The nutritional status and treatment of patients with hyperemesis gravidarum. Am J Obstet Gynecol 172, 1585–1591 (1995).

70. G. Koren, A. Ornoy, M. Berkovitch, Hyperemesis gravidarum-Is it a cause of abnormal fetal brain development? Reprod Toxicol 79, 84–88 (2018).

71. R. W. Smithells, S. Sheppard, C. J. Schorah, Vitamin deficiencies and neural tube defects. Arch Dis Child 51, 944–950 (1976).

72. A. L. Chandler et al., Neural tube defects and maternal intake of micronutrients related to one-carbon metabolism or antioxidant activity. Birth Defects Res A Clin Mol Teratol 94, 864–874 (2012).

73. M. V. S. R. Group, Prevention of neural tube defects: results of the Medical Research Council vitamin study. The Lancet 338, 131–137 (1991).

74. A. Rumbold, C. A. Crowther, Vitamin C supplementation in pregnancy. *Cochrane Database Syst Rev*, CD004072 (2005).

75. A. Rumbold, E. Ota, C. Nagata, S. Shahrook, C. A. Crowther, Vitamin C supplementation in pregnancy. Cochrane Database Syst Rev 2015, CD004072 (2015).

76. O. O. Balogun et al., Vitamin supplementation for preventing miscarriage. Cochrane Database Syst Rev 2016, CD004073 (2016).

77. M. Viswanathan et al., Folic acid supplementation for the prevention of neural tube defects: an updated evidence report and systematic review for the US Preventive Services Task Force. JAMA 317, 190–203 (2017).

78. MRC Vitamin Study Research Group, Prevention of neural tube defects: results of the Medical Research Council Vitamin Study. The Lancet 338, 131-137 (1991).

79. S. Yamaguchi et al., Tet1 controls meiosis by regulating meiotic gene expression. Nature 492, 443–447 (2012).

80. S. Yamaguchi, L. Shen, Y. Liu, D. Sendler, Y. Zhang, Role of Tet1 in erasure of genomic imprinting. Nature 504, 460–464 (2013).

81. K. Theiler, *The house mouse; development and normal stages from fertilization to 4 weeks of age*. (Springer-Verlag, Berlin, New York,, 1972), pp. 168 p.

82. M. Szultka, M. Buszewska-Forajta, R. Kaliszan, B. Buszewski, Determination of ascorbic acid and its degradation products by high-performance liquid chromatography-triple quadrupole mass spectrometry. ELECTROPHORESIS 35, 585–592 (2014).

83. H. Feng, H. Wu, Differential methylation analysis for bisulfite sequencing using DSS. Quant Biol 7, 327–334 (2019).

84. K. D. Hansen, B. Langmead, R. A. Irizarry, BSmooth: from whole genome bisulfite sequencing reads to differentially methylated regions. Genome Biol 13, R83 (2012).

85. A. Kechin, U. Boyarskikh, A. Kel, M. Filipenko, cutPrimers: A New Tool for Accurate Cutting of Primers from Reads of Targeted Next Generation Sequencing. J Comput Biol 24, 1138–1143 (2017).

86. A. Dobin et al., STAR: ultrafast universal RNA-seq aligner. Bioinformatics 29, 15–21 (2013).

87. T. Smith, A. Heger, I. Sudbery, UMI-tools: modeling sequencing errors in Unique Molecular Identifiers to improve quantification accuracy. Genome Res 27, 491–499 (2017).

88. Y. Liao, G. K. Smyth, W. Shi, featureCounts: an efficient general purpose program for assigning sequence reads to genomic features. Bioinformatics 30, 923–930 (2014).

89. B. Li, C. N. Dewey, RSEM: accurate transcript quantification from RNA-Seq data with or without a reference genome. BMC Bioinformatics 12, 323 (2011).

90. M. I. Love, W. Huber, S. Anders, Moderated estimation of fold change and dispersion for RNA-seq data with DESeq2. Genome Biol 15, 550 (2014).

91. M. Stephens, False discovery rates: a new deal. Biostatistics 18, 275–294 (2017).

92. G. Yu, L. G. Wang, Y. Han, Q. Y. He, clusterProfiler: an R package for comparing biological themes among gene clusters. OMICS 16, 284–287 (2012).

