## Supplementary Figures for "Maternal Vitamin C Deficiency and Genetic Risk Factors Contribute to Congenital Defects through Dysregulation of DNA Methylation"

### van der Veer et al. Supplementary Figure 1

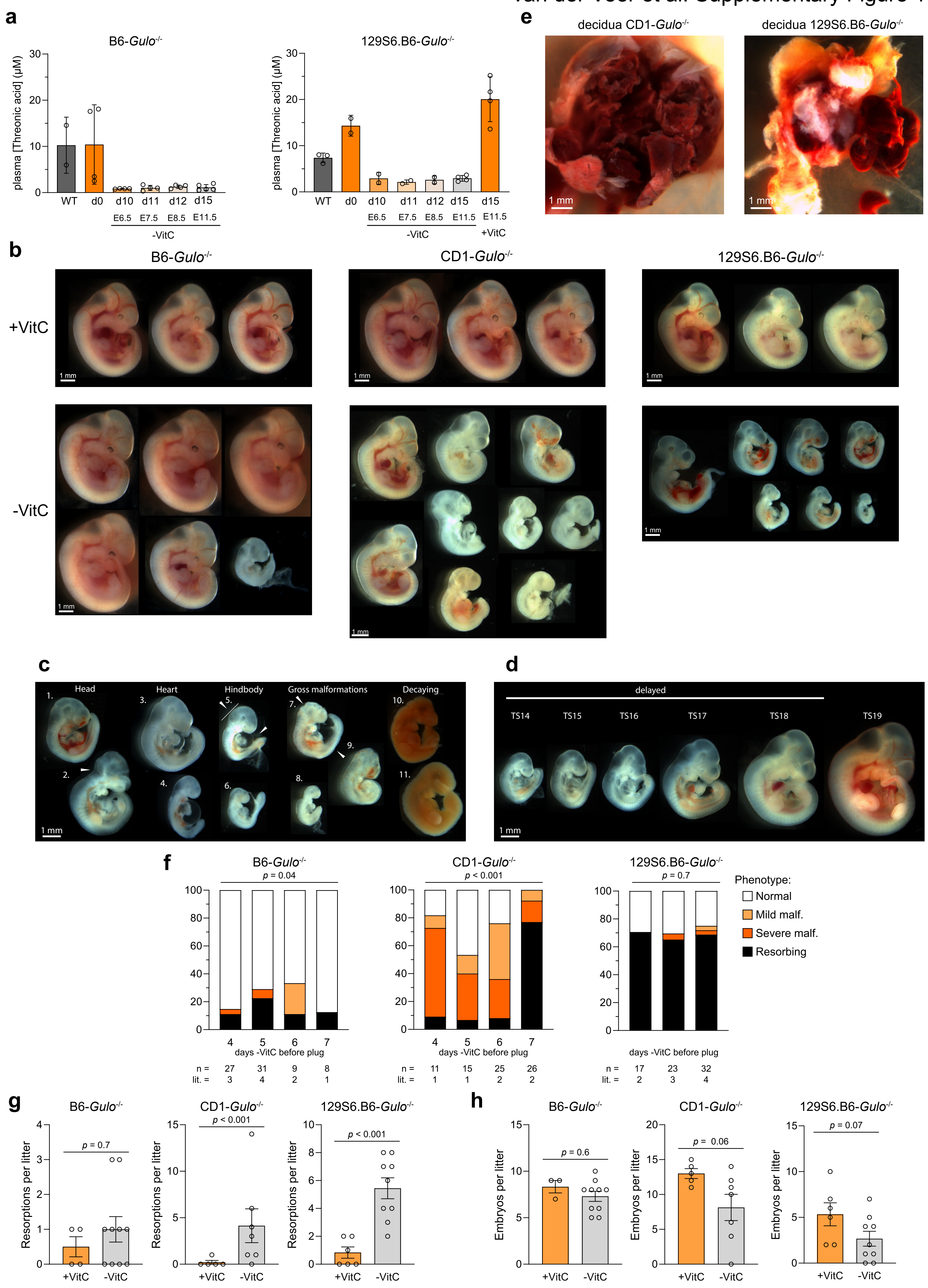

**Supplementary Figure 1 (related to main Figure 1)**

**a**, Plasma levels of total ascorbate detected as the degradation product threnoic acid by LC-MS/MS in B6-*Gulo*$^{-/-}$ and 129S6.B6*Gulo*$^{-/-}$ female mice at the indicated timepoints of VitC withdrawal, compared to WT mice per strain. d0, dam maintained on 1% stabilized VitC diet prior to VitC withdrawal. The gestational time corresponding to days of maternal VitC withdrawal assumes copulation within the first day of pairing with a stud male, i.e. day 4 after VitC withdrawal. **b**, Three representative +VitC *Gulo*$^{-/-}$ embryos and a representative full litter of -VitC *Gulo*$^{-/-}$ embryos of each background. **c**,**d**, Representative images of malformations (c) and delays (d) observed in VitC-deprived embryos. Overview images in (b-d) are assembled from individual embryo pictures. Scale bars indicate 1 mm. **e,** Representative images of hemorrhagic placentas in -VitC CD-*Gulo*$^{-/-}$ and 129S6.B6-*Gulo*$^{-/-}$. **f,** Phenotype penetrance scores of -VitC B6, CD1, and 129S6.B6 *Gulo*$^{-/-}$ embryos, stratified by days of VitC depletion before detection of copulation plug, i.e. fertilization. Chi-square test was used for statistics. "Mild malformation" (malf.) = 1 malformation, "severe malformation" = multiple malformations or grossly malformed. "Resorbing" = empty decidua or embryos in the process of being resorbed. "no TS" = not possible to assign a Theiler stage. n = number of embryos per condition, litters (lit.) = number of litters per condition. Statistical comparisons are made using Chi-square test. **g**, Number of resorptions per litter, scored as incidences of resorbing embryos in the process being resorbed or empty decidua. **h**, Number of embryos per genetic background and treatment. Pairwise comparisons are done using Mann-Whitney U test.

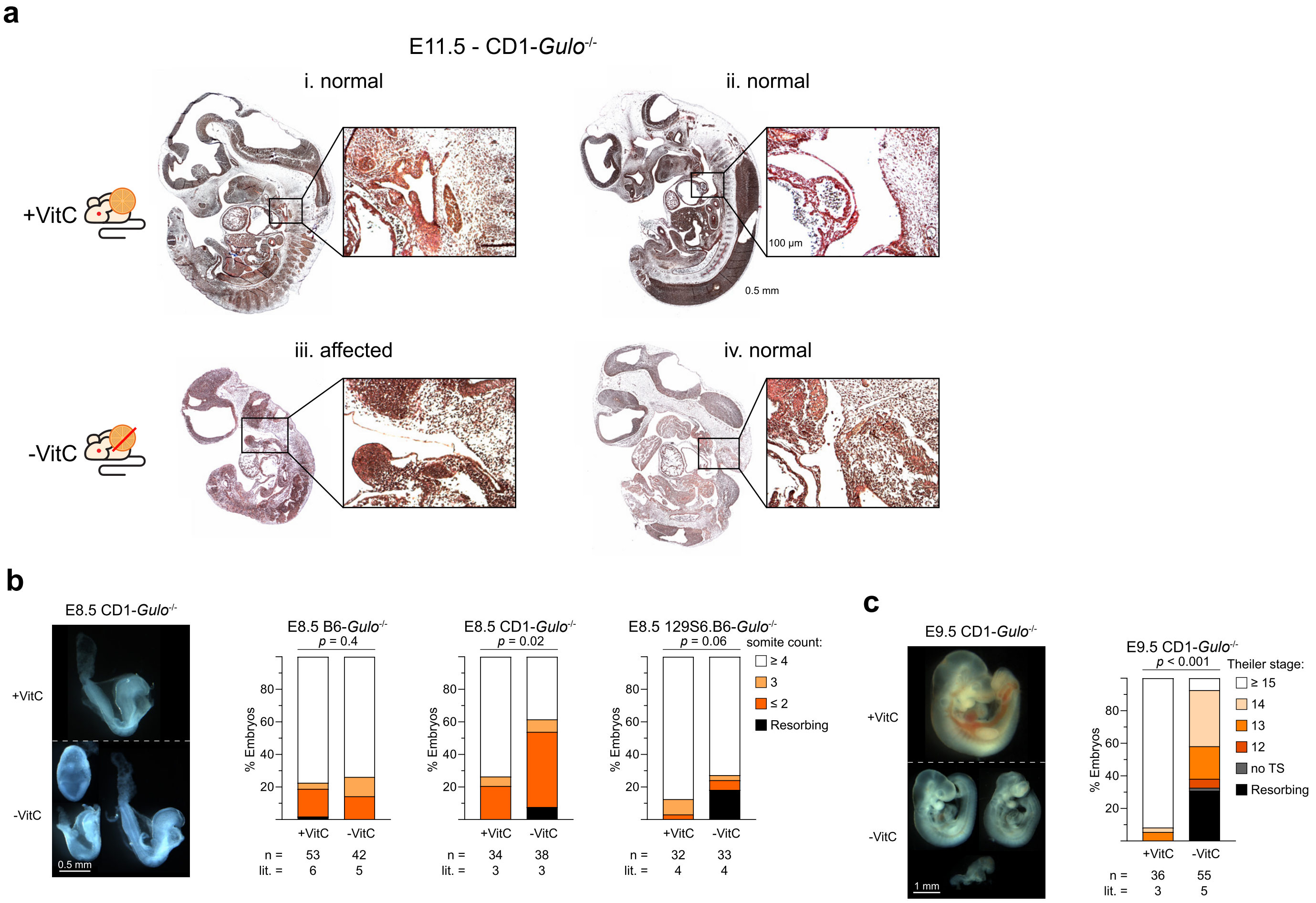


**Supplementary Figure 2 (related to main Figure 1)**

**a**, Sirius red staining (red = collagen) in +VitC (i, ii) and -VitC (iii, iv) CD1-*Gulo*$^{-/-}$ embryos. **i**, **ii**, **iv**, are embryos scored as "normal", while **iii** is an embryo with severe malformations "affected" by VitC deprivation. Scale bars in the insets indicate 100 μM, while scale bars in the whole embryo images indicate 0.5 mm. **b**, **c**, Representative images of E8.5 (b, left) and E9.5 (c, left) +VitC and -VitC *Gulo*$^{-/-}$ embryos and associated staging scores (right) in B6, CD1, and 129S6.B6 genetic backgrounds at E8.5 (b) and in CD1 background at E9.5 (**c**). The scale bar is 0.5 mm for E8.5 and 1 mm for E9.5 images. E8.5 embryos were staged by somite counts, not Theiler stage. "mild malformation" (malf.) = 1 malformation, "severe malformation" = multiple malformations or grossly malformed, "Resorbing" = empty decidua or embryos in the process of being resorbed, "no TS" = not possible to assign a Theiler stage, n = number of embryos per condition, litters (lit.) = number of litters per condition. Pairwise comparisons were performed using Chi-square test.

**a**

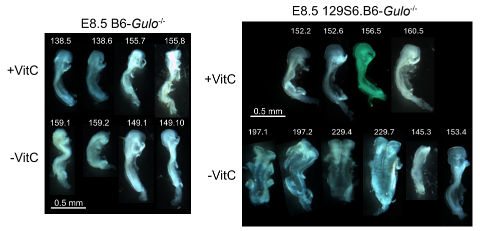


**c**

**d**

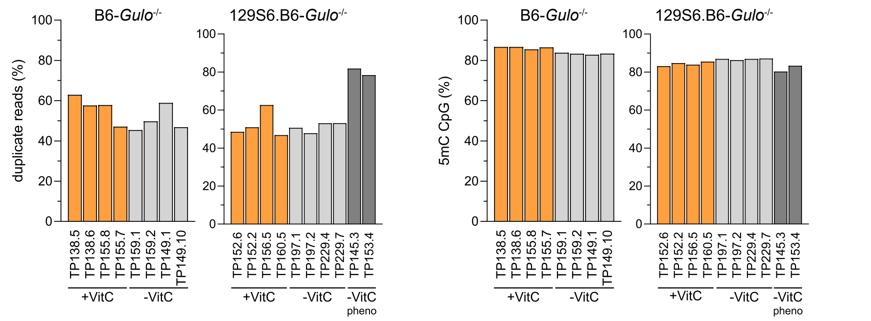


**b**

| Background | Condition | Sex | TP ID | Somite Count | Coverage Mean | Coverage Median | 5mC (%) CHG | 5mC (%) CHH |
|---|---|---|---|---|---|---|---|---|
| B6 | +VitC | ♂ | TP138.5 | 6 | 7,5 | 6 | 1,4 | 1,1 |
| | | ♀ | TP138.6 | 4 | 5,8 | 4 | 1,4 | 1,1 |
| | | ♂ | TP155.8 | 6 | 7,6 | 7 | 1,3 | 1,0 |
| | | ♀ | TP155.7 | 5 | 8,3 | 7 | 1,3 | 1,0 |
| | -VitC | ♂ | TP159.1 | 5 | 9,5 | 9 | 1,4 | 1,0 |
| | | ♀ | TP159.2 | 4 | 9,7 | 9 | 1,4 | 1,0 |
| | | ♂ | TP149.1 | 6 | 7,9 | 7 | 1,3 | 1,0 |
| | | ♀ | TP149.10 | 6 | 8,6 | 8 | 1,5 | 1,1 |
| 129S6.B6 | +VitC | ♂ | TP152.6 | 6 | 6,4 | 5 | 1,5 | 1,1 |
| | | ♀ | TP152.2 | 5 | 6,4 | 5 | 1,5 | 1,1 |
| | | ♂ | TP156.5 | 6 | 6,5 | 5 | 1,5 | 1,1 |
| | | ♀ | TP160.5 | 5 | 5,3 | 4 | 1,6 | 1,3 |
| | -VitC | ♂ | TP197.1 | 5 | 3,9 | 3 | 1,7 | 1,3 |
| | | ♀ | TP197.2 | 5 | 4,9 | 4 | 1,7 | 1,3 |
| | | ♂ | TP229.4 | 5 | 7,5 | 5 | 1,5 | 1,1 |
| | | ♂ | TP229.7 | 5 | 7,5 | 5 | 1,8 | 1,3 |
| | -VitC pheno | ♂ | TP145.3 | 4 | 2,4 | 2 | 1,3 | 1,1 |
| | | ♀ | TP153.4 | 6 | 3,8 | 3 | 1,4 | 1,0 |

-VitC

**e**

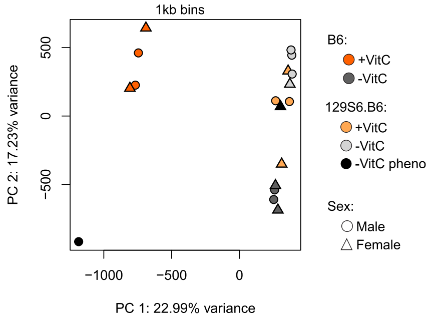


**f**

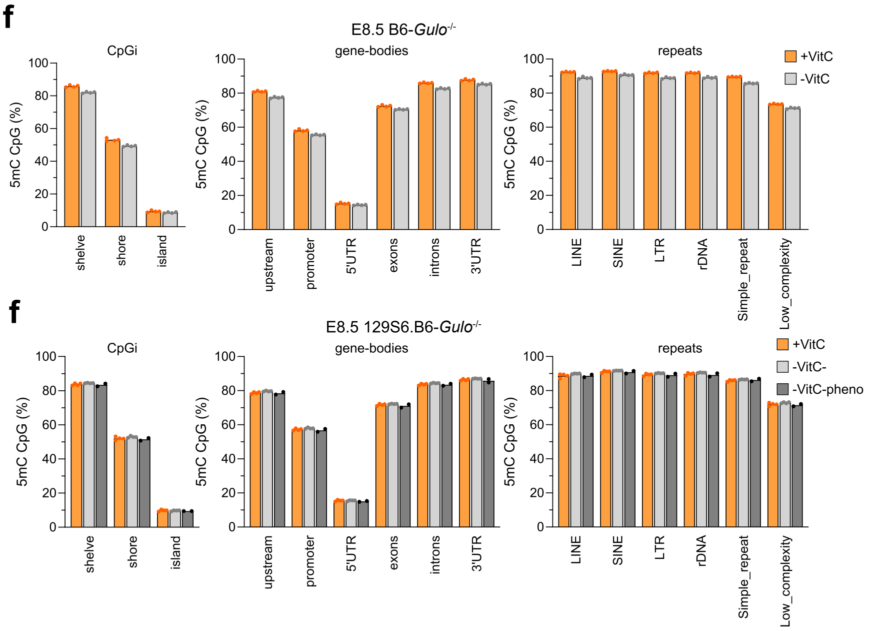


**h**

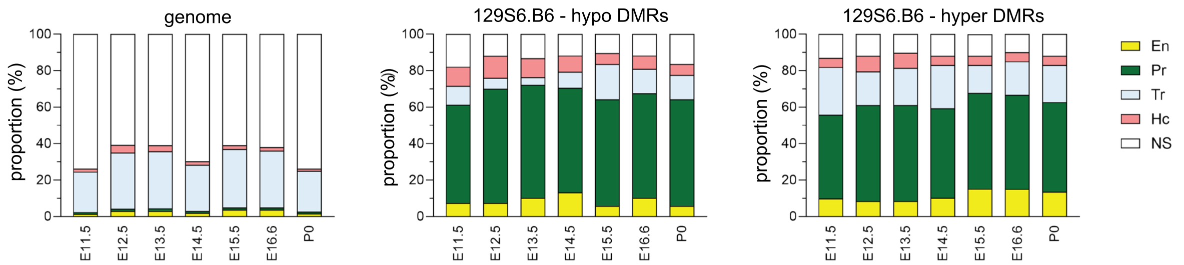


**i**

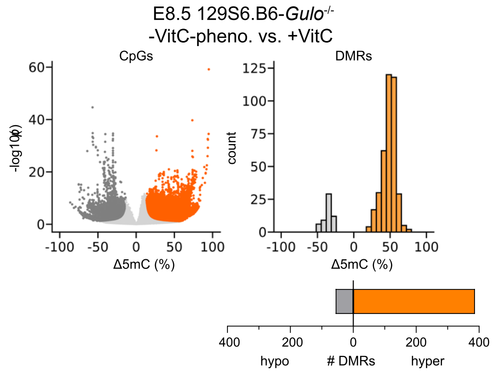


**j**

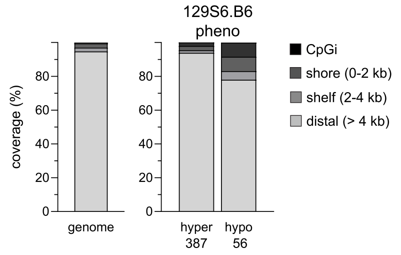

**Supplementary Figure 3 (related to main Figure 2)**

**a**, Images of the individual E8.5 embryos collected for WGBS. **b**, Table listing the experimental identifiers, sex, somite counts of individual embryos used in WGBS and the sample library coverage and global non-CpG methylation levels. **c**, **d**, Duplication rate (**c**) and global CpG methylation levels (**d**) of WGBS libraries. **e**, PCA of CpG methylomes in all samples grouped by strains (B6 and 129S6.B6), treatment (-VitC and +VitC) and sex (male and female). The two 129S6.B6 -VitC pheno samples are separately shown in black color. **f, g,** Global CpG methylation levels at indicated genomic elements per library from *Gulo*$^{-/-}$ headfolds of B6 (f) and 129S6.B6 (g) strains. **h**, Enrichment of 129S6.B6 DMRs overlapping with functional genomic elements active during mouse fetal development (timepoints are indicated) as defined in ENCODE3. The 15 chromatin states shown in Figure 2h condensed into 5 classes: En, enhancers; Pr, promoters; Tr, transcription; Hc, heterochromatin and NS, no signal. **i**, Volcano plot of differentially methylated CpGs, highlighted in orange and grey indicating those with significant gain or loss, respectively, in -VitC relative to +VitC, (left), histogram of DMR distribution by differences in CpG methylation levels in intervals of 10% (right), and number of hyper and hypo DMRs (bottom) in the two outlier 129S6.B6-*Gulo*$^{-/-}$ -VitC pheno headfolds compared to +VitC controls (n=4). **j**, Distribution of hyper- and hypo- DMRs by proximity to CpG islands (CpGi) in the two 129S6.B6-*Gulo*$^{-/-}$ -VitC pheno headfolds.

**a**

| Background | Condition | Sex | TP ID | Somite Count | Exclude |
|---|---|---|---|---|---|
| B6 | +VitC | ♂ | TP154.7 | 5 | |
| | | ♀ | TP155.5 | 5 | yes |
| | | ♂ | TP158.4 | 6 | |
| | | ♂ | TP158.5 | 6 | |
| | | ♀ | TP158.6 | 6 | |
| | | ♀ | TP158.7 | 5 | |
| | -VitC | ♂ | TP140.1 | 4 | |
| | | ♂ | TP140.2 | 5 | |
| | | ♂ | TP149.6 | 6 | |
| | | ♀ | TP149.9 | 6 | |
| | | ♀ | TP159.3 | 4 | |
| | | ♀ | TP159.4 | 4 | |
| | | ♂ | TP159.5 | 5 | |
| | | ♀ | TP159.7 | 4 | |
| 129S6.B6 | +VitC | ♂ | TP114.2 | 6 | |
| | | ♂ | TP114.5 | 6 | |
| | | ♀ | TP152.4 | 5 | |
| | | ♂ | TP152.5 | 6 | |
| | | ♀ | TP156.6 | 6 | |
| | | ♀ | TP160.1 | 6 | |
| | | ♂ | TP160.2 | 6 | |
| | | ♀ | TP160.6 | 6 | |
| | -VitC | ♂ | TP145.6 | 5 | |
| | | ♂ | TP153.5 | 4 | |
| | | ♀ | TP153.6 | 5 | |
| | | ♀ | TP197.3 | 6 | |
| | | ♂ | TP197.4 | 6 | |
| | | ♀ | TP198.2 | 5 | |
| | | ♂ | TP198.4 | 6 | |

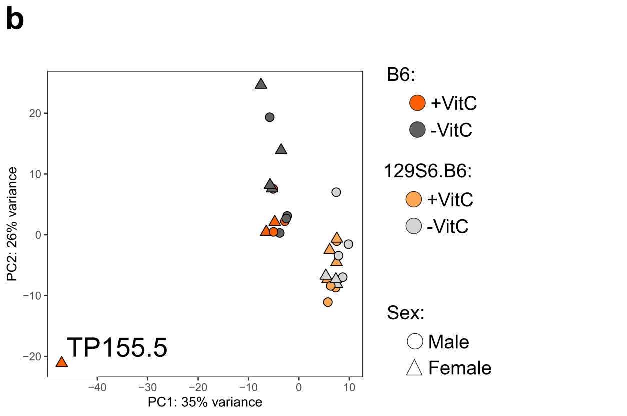


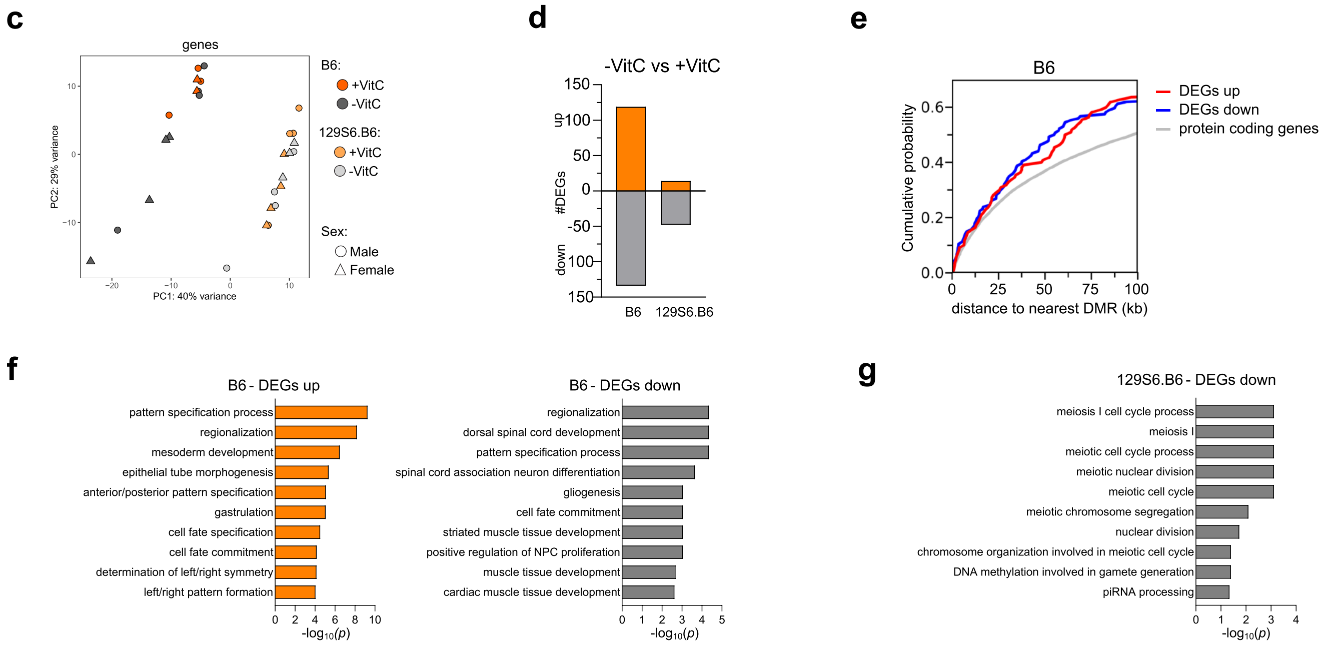


**Supplementary Figure 4 (related to main Figure 2)**

**a**, Table listing the experimental identifiers, sex and somite counts of E8.5 embryos used in RNA-sequencing. **b**, PCA plot of all RNA-seq data from all samples showing an outlier TP155.5 (B6, +VitC, female). **c**, PCA plot of transcriptomes after exclusion of outlier TP155.5. **d**, Number of RNA-seq DEGs defined by pairwise comparison of -VitC versus +VitC treatments per *Gulo*$^{-/-}$ strain, classified by up or downregulation relative to +VitC. **e**, Cumulative distribution of the distance to the nearest B6 hypo-DMR among up- and down-regulated DEGs in B6-*Gulo*$^{-/-}$ headfolds. **f,** Top 10 GO terms associated with up- (left) and down-regulated (right) DEGs in E8.5 B6-*Gulo*$^{-/-}$ headfolds. **g**, Top 10 GO terms associated with down-regulated DEGs in 129S6.B6-*Gulo*$^{-/-}$ headfolds.

van der Veer et al. Supplementary figure 5

a

| Background | Sex | [VitC] mg/ml | Phenotype | TP ID | TS | Coverage Mean | Coverage Median | 5mC (%) CHG | 5mC (%) CHH |
|---|---|---|---|---|---|---|---|---|---|
| CD1 | ♂ | 3300 | normal | TP178.9 | 19 | 22,8 | 23 | 1,1 | 1 |
| | | 330 | normal | TP185.10 | 19 | 19,9 | 20 | 1,1 | 1,1 |
| | | 100 | normal | TP199.1 | 19 | 14,0 | 14 | 1 | 1 |
| | | | NTD | TP199.3 | 19 | 12,9 | 13 | 1 | 1 |
| | | 33 | normal | TP190.3 | 19 | 22,1 | 22 | 1,2 | 1,1 |
| | | | delay | TP190.8 | x | 19,5 | 20 | 1,1 | 1,1 |
| | | 0 | delay | TP181.6 | x | 11,9 | 11 | 1,2 | 1,2 |
| | | rescue A6.5 | normal | TP72.5 | 19 | 13,0 | 13 | 1,1 | 1 |

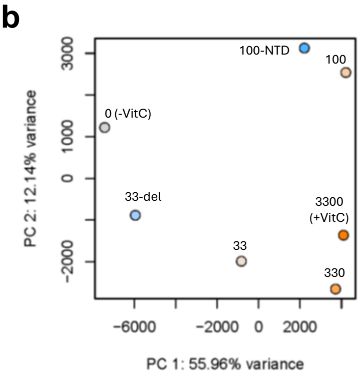


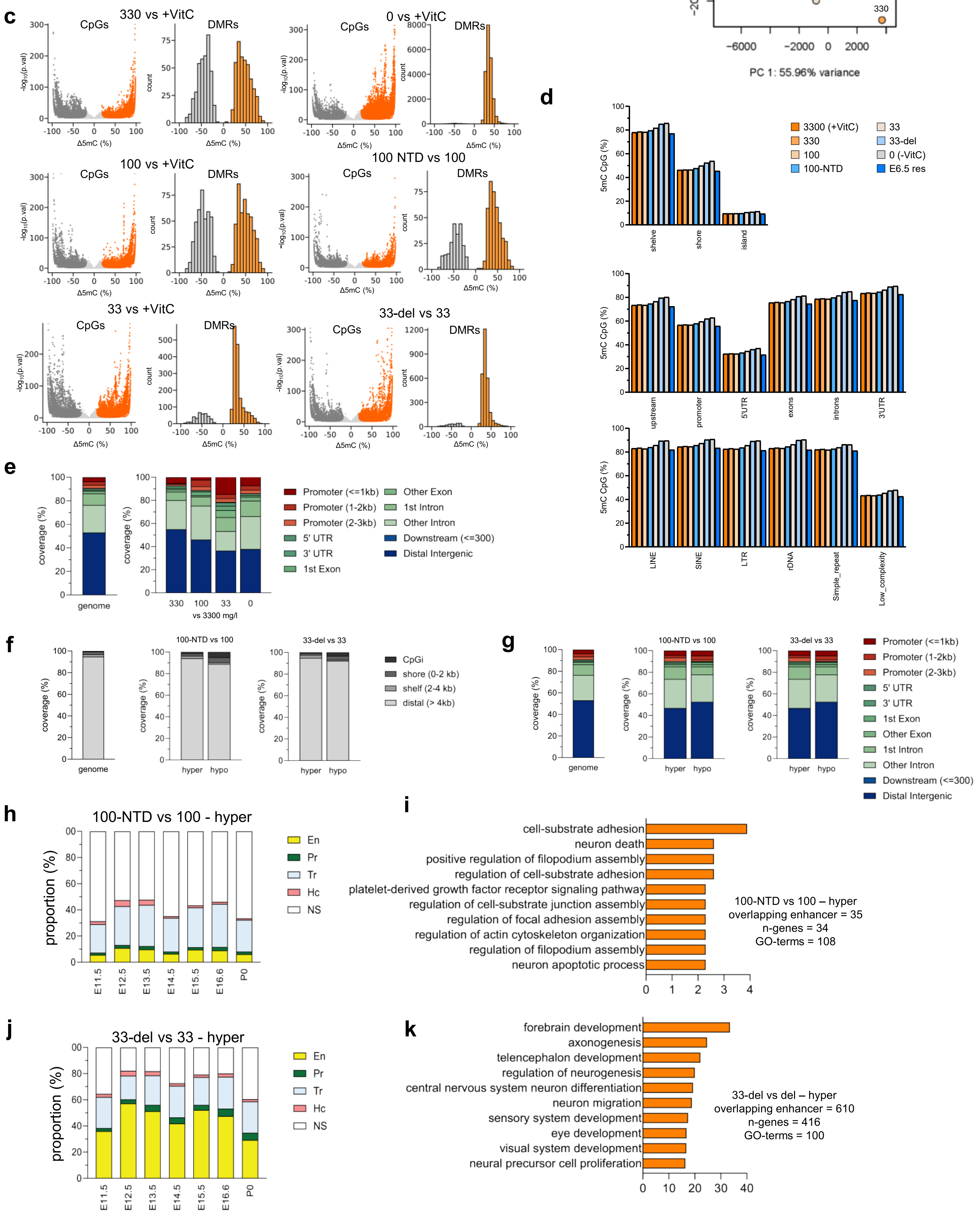

**Supplementary Figure 5 (related to main Figure 3)**

**a**, Table listing the experimental identifiers, phenotypes, developmental stages of the individual embryos used for WGBS and the sample library coverage and global non-CpG methylation levels. **b**, PCA plot of WGBS data from E11.5 brains, collected from each VitC dose exposure from 3300-0 mg/l. **c**, Volcano plots of differentially methylated CpGs, highlighted in orange and grey to indicate those with significant gain or loss, respectively, and histogram of DMR distribution by differences in CpG methylation levels in intervals of 10% (right) in pairwise comparisons of morphologically normal E11.5 brains from each VitC dose exposure at 330 ml/l, 100 mg/l and 33 mg/l compared to +VitC (3300 mg/l) control (left column), in completely VitC-deprived and deformed (0 mg/l, -VitC) compared to +VitC control brains and in phenotypically affected brains compared to morphologically normal brains at 100 mg/l and 33 mg/l (right column). **d**, Global CpG methylation levels within genomic features associated with CpGi, gene bodies and retrotransposable repeats across the genome in staged-matched morphologically normal E11.5 brains exposed to dose-titration of maternal VitC (orange to grey colouring) and those with phenotypes (blue). **e**, Distribution of DMRs induced by half logarithmic dose reduction in VitC until 33 mg/l in morphologically normal E11.5 brains and in completely VitC deprived brain, per pairwise comparison relative to +VitC control, at gene body elements. **f**, **g**, Distribution of hyperDMRs and hypoDMRs associated with NTD or growth delay versus morphologically normal brains per VitC dose by CpG island proximity (f) and genetic elements. **h-i**, Enrichment of NTD-associated hyper-DMRs at 100 mg/l VitC treatment at ENCODE3 genomic elements active during mouse fetal development from E11.5 to P0 (h) and the top 10 enriched GO terms overlapping with ENCODE3 enhancers (i). **j**, **k**, Enrichment of growth delay-associated hyper-DMRs at 33 mg/l VitC treatment at ENCODE3 genomic elements (j) and the top 10 enriched GO terms overlapping with ENCODE3 enhancers (k). Refer to genome reference of ENCODE3 functional elements and legends in Supplementary Figure 3h.

**a**

| Background | [VitC] mg/ml | Phenotype | Sex | TP ID | Theiler stage |
|---|---|---|---|---|---|
| CD1 | 3300 | normal | ♂ | TP178.3 | 19 |
| | | | ♀ | TP178.7 | 19 |
| | 330 | normal | ♂ | TP185.3 | 19 |
| | | | ♀ | TP185.7 | 19 |
| | 100 | normal | ♂ | TP199.15 | 19 |
| | | | ♀ | TP203.4 | 19 |
| | | NTD | ♂ | TP203.5 | 19 |
| | | | ♀ | TP199.14 | 19 |
| | 33 | normal | ♂ | TP190.4 | 19 |
| | | | ♀ | TP190.10 | 19 |
| | rescue E6.5 | normal | ♂ | TP72.9 | 19 |
| | | | ♀ | TP72.10 | 19 |

**b**

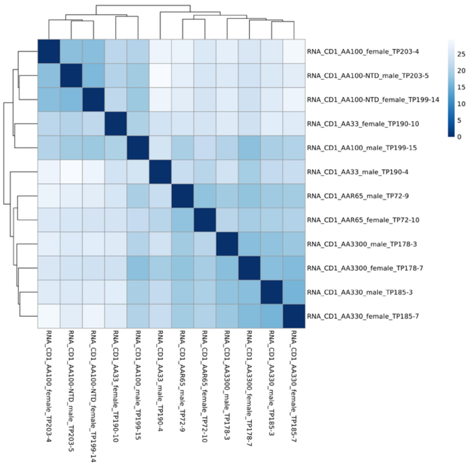

**c**

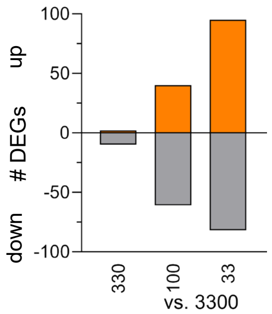


**d**

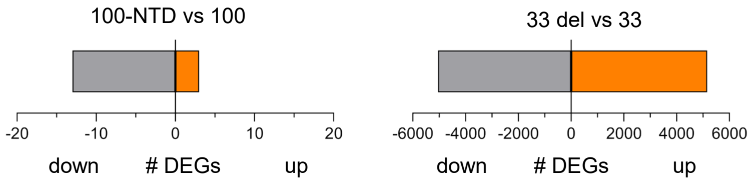


**Supplementary Figure 6 (related to main Figure 3)**

**a**, Table listing the experimental identifiers, sex, phenotype and stage of the individual E11.5 CD1-*Gulo*$^{-/-}$ embryos used for RNA-sequencing. **b**, Heatmap with hierarchical clustering based on sample-to-sample distances of samples with VitC dose treatment from 3300-0 mg/l and rescue at E6.5. **c**, Number of RNA-seq DEGs defined by pairwise comparison of each indicated VitC dose treatment vs +VitC, classified by up or downregulation. **d**, Number of RNA-seq DEGs defined by pairwise comparison of 100 mg/l -NTD vs 100 mg/l (left) and 33 mg/l -del vs 33 mg/l (right), classified by up or downregulation.

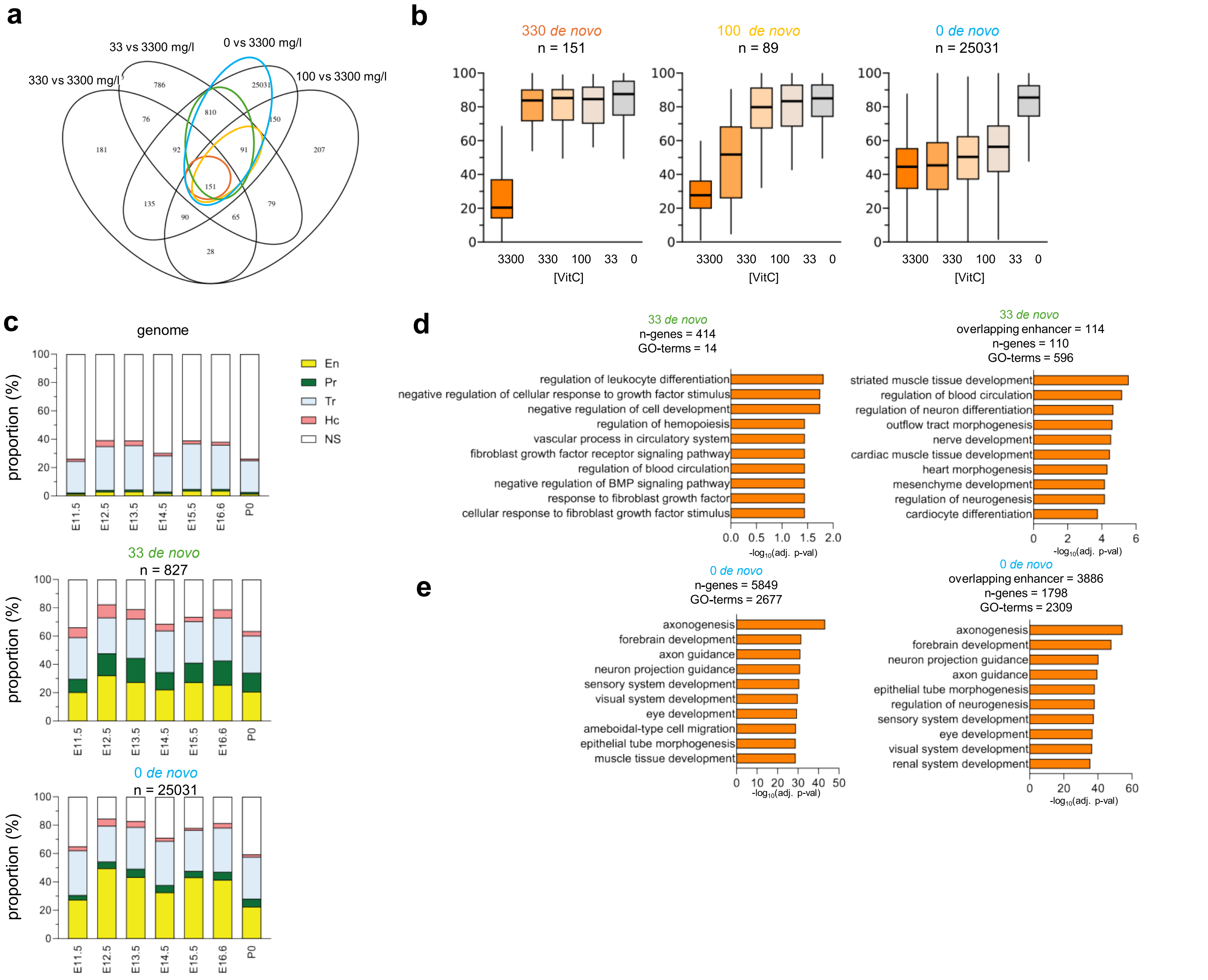


**Supplementary Figure 7 (related to main Figure 3)**

**a**, Venn diagram showing overlap of hyperDMRs from all pairwise comparisons of each VitC dose treatment versus +VitC control to define *de novo* hyperDMRs (circled in color) that accrue with each half logarithmic dose reduction and stayed hypermethylated at subsequently lower doses. **b**, Dose-response of CpG methylation levels in subsets of *de novo* hyperDMRs induced by each titrated VitC dose reduction from 3300 to 0 mg/l. The box represents the interquartile range (IQR) and the line within indicates the median. **c**, Enrichment of *de novo* hyper-DMRs at 33 mg/l and 0 mg/l VitC dose treatments at ENCODE3 genomic elements active throughout mouse fetal development. **d**, Top 10 enriched GO terms of 33 mg/l *de novo* hyperDMRs that are overlapping with ENCODE3 enhancers. **e**, Top 10 enriched GO terms of 0 mg/l *de novo* hyperDMRs that are overlapping with ENCODE3 enhancers.

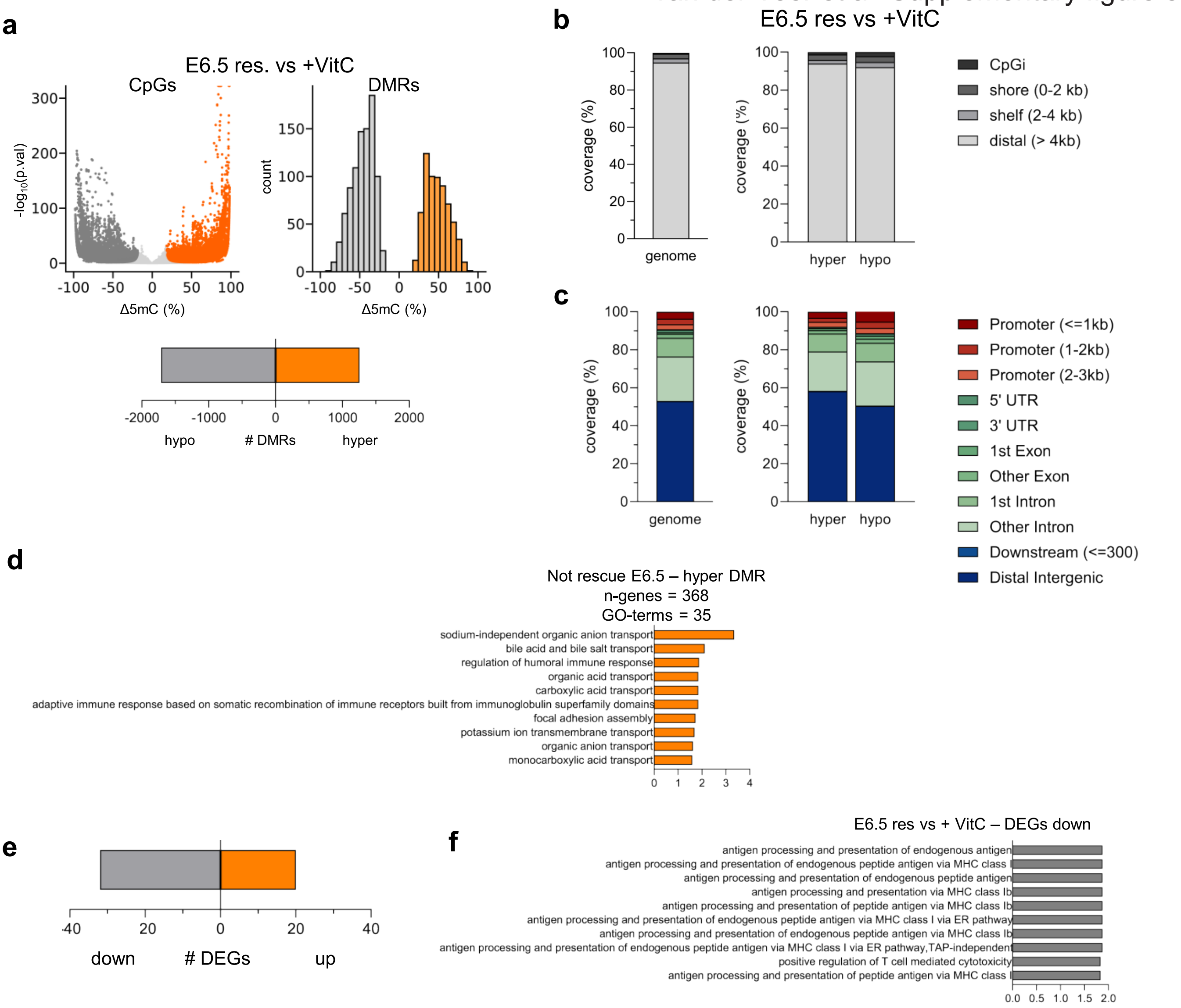


**Supplementary Figure 8 (related to main Figure 4)**

**a**, Volcano plot of differentially methylated CpGs, highlighted in orange and grey to indicate those with significant gain or loss, respectively, in an E11.5 CD1-*Gulo*$^{-/-}$ brain fully rescued by high dose VitC re-supplementation at E6.5 relative to +VitC brain (left), histogram of DMR distribution by differences in CpG methylation levels in intervals of 10% (right), and number of hyper and hypo DMRs (bottom) in **b**, **c,** Distribution of hyperDMRs and hypoDMRs by CpG island proximity (b) and gene body elements (c) in E11.5 brains after re-supplementation at E6.5. **d**, Top 10 enriched GO terms of resistant hyper hyperDMRs resistant to VitC re-supplementatio, as defined in Figure 4f. **e,** Number of RNA-seq DEGs defined by pairwise comparison of a VitC-deprived E11.5 brains (n=2) rescued at E6.5 vs VitC+ control brains (n=2). **f**, Top 10 GO terms associated down-regulated DEGs defined by pairwise comparison of VitC-deprived brains rescued at E6.5 vs VitC+ controls.

van der Veer et al. Supplementary figure 9

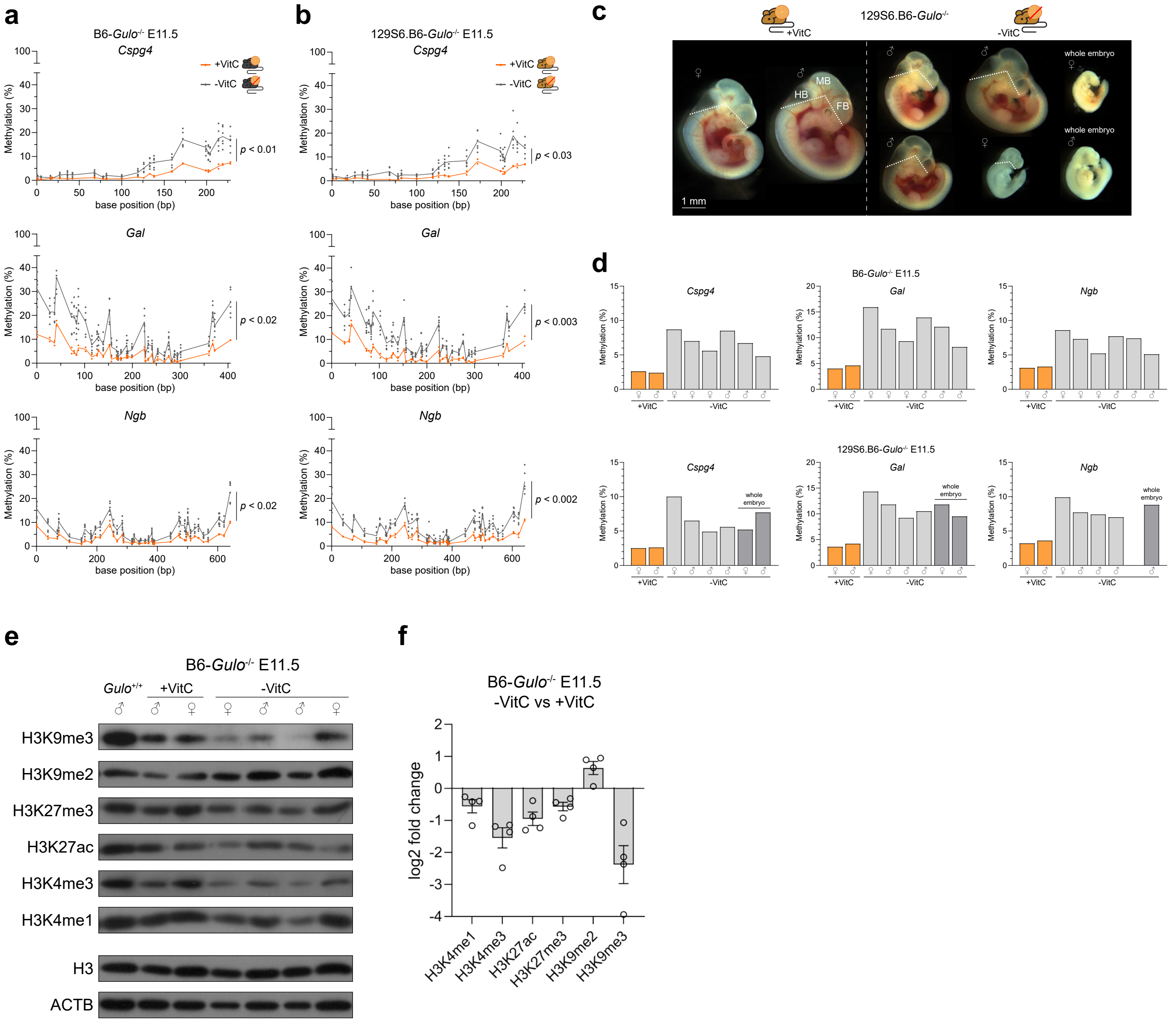


#### Supplementary Figure 9

**a**, **b**, CpG methylation levels at the promoters of *Cspg4*, *Gal*, and *Ngb* in E11.5 brains collected from +VitC and -VitC B6 (**a**) and 129S6.B6 (**b**) *Gulo*$^{-/-}$ embryos. n = 2 individual embryos for +VitC, n = 6 individual embryos for -VitC for both B6 and 129S6.B6. Pairwise statistical comparison is made using a paired t-test. **c**, Images of actual embryos used for targeted bisulfite sequencing in **a**, **b**, from +VitC and -VitC 129S6.B6-*Gulo*$^{-/-}$ embryos. We collected the entire brain, consisting of the hindbrain (HB), midbrain (MB), and forebrain (FB). Dotted lines indicate dissection planes. The image is assembled from individual embryo pictures. **d**, Non-weighted average CpG methylation per amplicon, per individual E11.5 brain collected from +VitC and -VitC B6 and 129S6.B6-Gulo-/- embryos. Dark grey denotes samples in which whole embryos were collected because of gross malformations. **e**, Western blot showing global H3K9me3, H3K9me2, H3K27me3, H3K27ac, H3K4me3, H3K4me1 content in E11.5 brains collected from +VitC and -VitC B6-*Gulo*$^{-/-}$ embryos. As a control, a B6-*Gulo*$^{+/+}$ is also included. H3 and ACTB are used as loading controls. **f**, ImageJ densitometric measurement of each histone mark in -VitC samples relative to VitC-replete controls.
